# Resource uptake and the evolution of moderately efficient enzymes

**DOI:** 10.1101/2020.11.08.373290

**Authors:** Florian Labourel, Etienne Rajon

## Abstract

Enzymes speed up reactions that would otherwise be too slow to sustain the metabolism of self-replicators. Yet, most enzymes seem only moderately efficient, exhibiting kinetic parameters orders of magnitude lower than their expected physically achievable maxima and spanning over surprisingly large ranges of values. Here, we question how these parameters evolve using a mechanistic model where enzyme efficiency is a key component of individual competition for resources. We show that kinetic parameters are under strong directional selection only up to a point, above which enzymes appear to evolve under near-neutrality, thereby confirming the qualitative observation of other modeling approaches. While the existence of a large fitness plateau could potentially explain the extensive variation in enzyme features reported, we show using a population genetics model that such a widespread distribution is an unlikely outcome of evolution on a common landscape, as mutation-selection-drift balance occupy a narrow area even when very moderate biases towards lower efficiency are considered. Instead, differences in the evolutionary context encountered by each enzyme should be involved, such that each evolves on an individual, unique landscape. Our results point to drift and effective population size playing an important role, along with the kinetics of nutrient transporters, the tolerance to high concentrations of intermediate metabolites, and the reversibility of reactions. Enzyme concentration also shapes selection on kinetic parameters, but we show that the joint evolution of concentration and efficiency does not yield extensive variance in evolutionary outcomes when documented costs to protein expression are applied.

## 1 Introduction

Living organisms need to uptake and metabolize nutrients, relying on enzymes to catalyse chemical reactions along metabolic pathways. Accordingly, and despite being intrinsically reversible (Haldane, 1930; Klipp and Heinrich, 1994), *in vivo* enzyme-catalyzed reactions are commonly thought of as an irreversible two-step process (Bar-Even *et al.*, 2011, 2015; Michaelis and Menten, 1913; Johnson and Goody, 2011):

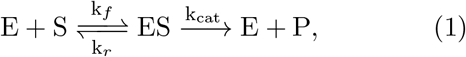

where *k_f_* and *k_r_* are the rates of association and dissociation between enzyme and substrate, and *k*_cat_ is the turnover number, that is the rate of formation of the product *P* from *ES* complexes. The first part of this chemical equation describes the encounters between the enzyme *E* and the substrate *S*; the enzyme will be efficient if *ES* complexes form often and do not dissociate before the substrate has been turned into a product, which is reflected by the constant 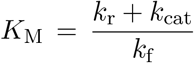. The efficiency *v* of an enzyme – the rate at which it makes a product *P* from *S* – depends on these two constants through equation (2):

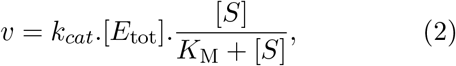

under the assumption that the concentration [*S*] is approximately constant and that of [*ES*] is at steady-state (Michaelis and Menten, 1913; Briggs and Haldane, 1925).

At first glance, Natural Selection is presumed to optimize enzymatic efficiency by pushing *k*_cat_ upwards and *K*_M_ downwards to universal physical limits. Enzyme efficiencies are for instance limited by the diffusion properties of their aqueous environment, which sets an upper bound of approximately 10^8^ − 10^10^*M* ^−1^*s*^−1^ for the ratio *k*_cat_/*K*_M_ (Alberty and Hammes, 1958; Zhou and Zhong, 1982). Nearly optimal enzymes indeed seem to exist, as exemplified by triosephosphate isomerase (TIM) whose ratio is close to this theoretical ceiling (Knowles and Albery, 1977). But they are uncommon: most enzymes appear to be only moderately efficient and far off these physical limits – including enzymes immediately flanking TIM in the glycolysis metabolic pathway (Davidi *et al.*, 2018). Indeed, Bar-Even *et al.* (2011) have analysed a dataset of several hundreds of enzymes and found a wide diversity among enzyme parameters, thus sketching a puzzling pattern that has far more in common with a zoo than it looks like variations around an archetypal form.

This wide distribution of enzyme features could partly be explained by differences between enzyme behaviour *in vivo* and *in vitro*. Such differences are expected, first because diffusion in a test tube is hardly comparable to diffusion in the cytoplasm (Ellis, 2001; Rivas *et al.*, 2004; Zhou *et al.*, 2008; Rivas and Minton, 2018). As the cytoplasm gets packed, the cell approaches a state where molecules are less mobile, hindering substrate-enzyme encounters (Muramatsu and Minton, 1988; Zimmerman and Minton, 1993; Blanco *et al.*, 2018). In this regard, *K*_M_ values are likely underestimated *in vitro*, and enzymes should perform less efficiently *in vivo*. Simultaneously, macromolecular crowding can sometimes improve catalytic activity *in vivo*, making specificity constants *k*_cat_/*K*_M_ higher than their *in vitro* estimates (Ralston, 1990; Ellis, 2001; Jiang and Guo, 2007; Pozdnyakova and Wittung-Stafshede, 2010). Crowding effects are obviously important for our understanding of enzyme evolution but, alone, they are definitely too weak to explain the wide variability across enzymes insofar as their reported estimates typically lie in the range of one order of magnitude (Davidi *et al.*, 2016).

Another source of explanation to the observed distribution of enzymes (in)efficiencies is a failure of Evolution to consistently optimize them, possibly due to physical constraints. Indeed, Heckmann *et al.* (2018) have shown how a variety of *k*_cat_s may evolve provided that some of them are physically constrained. Besides the diffusion limit already mentioned, constraints on enzyme evolution might include an intrinsic trade-off (Gudelj *et al.*, 2010; Stiffler *et al.*, 2015) that originates from the dependency of both *k*_cat_ and *K*_M_ on intermediate energy profiles (Heinrich *et al.*, 1991). Nonetheless, this trade-off is scarcely observable among evolved enzymes – Bar-Even *et al.* (2011) report a coefficient of determination around 0.09 for the correlation between log_10_(*k*_cat_) and log_10_(*K*_M_) – suggesting that it can be overcome. Other constraints may exist and be specific of a given reaction (Klipp and Heinrich, 1994) – *e.g.* reaction reversibility – potentially explaining a part of the variance in enzyme efficiencies. It remains that estimating constraints on all individual enzymes appears like a daunting task, which could be guided, in part, by the identification of deviations from evolutionary predictions.

Following this idea, the premise of our theoretical investigation into the origins of enzyme diversity is that it results mainly from unconstrained evolution, such that the reported differences may be caused by the combined action of selection and genetic drift. It is important to notice that the information we have is partial, as an enzyme’s activity is the joint result of its kinetic constants and cellular concentration, perhaps also contributing to the reported variance in the former. In fact, Davidi *et al.* (2016)’s method to determine *in vivo* turn-over rates lends some credence to the idea that increased levels of expression make up for lower kinetic constants (Davidi *et al.*, 2018). It is therefore obvious that an enzyme’s expression needs to be considered as another dimension of its activity, especially since it has been shown that the evolutionary tuning of gene expression can happen very quickly (Dekel and Alon, 2005).

Concomitantly, an enzyme’s activity can be impacted by protein misfolding, which reduces the effective enzyme concentration (Tokuriki and Tawfik, 2009; Yue *et al.*, 2005; Drummond *et al.*, 2005; Echave and Wilke, 2017) while also impacting fitness by enhancing protein erroneous interactions (Yang *et al.*, 2012) and the formation of toxic protein aggregates (Bucciantini *et al.*, 2002; Sabate *et al.*, 2010; Geiler-Samerotte *et al.*, 2011). Protein stability is thus under strong purifying selection to avoid the deleterious effects of misfolding (Drummond and Wilke, 2008). Accordingly, it has been shown that proteins have evolved to stand beyond a stability threshold (Bloom *et al.*, 2005), although marginally (Taverna and Goldstein, 2002).

Because mutations are on average destabilizing, this definitely narrows down the spectrum of adaptive mutations (Shoichet *et al.*, 1995; DePristo *et al.*, 2005; Weinreich *et al.*, 2006; Tokuriki *et al.*, 2007, 2008; Lunzer *et al.*, 2010). Nevertheless, several studies have reported the existence of a genotype space where activity can be optimized without compromising stability (Schreiber *et al.*, 1994; van den Burg and Eijsink, 2002; Bloom *et al.*, 2004; Knies *et al.*, 2017; Miller, 2017). Even when improving function requires the fixation of destabilizing mutations, compensatory mutations can in principle cancel out stability losses arising from active site evolution (DePristo *et al.*, 2005; Tokuriki *et al.*, 2008; Tokuriki and Tawfik, 2009; Storz, 2018). Adaptive evolution may even be facilitated by preexisting mutational robustness against misfolding (Bloom *et al.*, 2006, 2007). Therefore, although the requirement of a stable, correctly folding protein may sometimes slow down the evolutionary process, it is rather unlikely that stability explains the distribution of enzyme kinetic parameters albeit marginally.

Enzyme kinetics evolution has often been considered theoretically through the lens of flux control (Burns *et al.*, 1985; Clark, 1991; Fell, 1992; Kacser *et al.*, 1995; Yi and Dean, 2019). Indeed, the control of the flux in a metabolic pathway is shared between all enzymes, in what is known as the summation theorem (Kacser and Burns, 1973; Heinrich and Rapoport, 1974). Thence, because the sum of control coefficients must equal 1 within a pathway, if all enzymes have similar kinetic parameters, none of them exerts a strong influence (Dean, 1995). But if one enzyme departs from this trend and becomes inefficient, it exerts a strong control at the expense of others (Dykhuizen and Dean, 1990). This leads to diminishing-returns epistasis in which the fitness landscape flattens because, as an enzyme becomes more efficient, subsequent mutations have smaller effects (Kacser and Burns, 1973; Dykhuizen *et al.*, 1987; Tokuriki *et al.*, 2012), a finding that has since received empirical confirmation (Fell, 1992; Dean, 1995; Lunzer *et al.*, 2005; Yi and Dean, 2019; Chou *et al.*, 2014).

Hartl *et al.* (1985) and Dean *et al.* (1986) have considered such a fitness landscape under a population genetics framework to conclude that enzymes may quickly reach a fitness plateau and evolve on nearly neutral landscapes (Ohta, 1992). Nonetheless, these studies fall short of explaining why inefficient enzymes having stronger control do not evolve higher activities (Yi and Dean, 2019). In these models as in most, an enzyme’s efficiency is captured by its activity, generally represented by a composite of *k*_cat_, *K*_M_ and enzyme concentration (Hartl *et al.*, 1985; Clark, 1991; Chou *et al.*, 2014; Kaltenbach and Tokuriki, 2014), such that the distinct evolutionary dynamics of these parameters, together with an enzyme’s concentration, is ignored. This reduction of an enzyme’s dimensionality goes against the empirical observation that each dimension may have a differential impact on fitness in the context of antibiotic resistance (Walkiewicz *et al.*, 2012; Stiffler *et al.*, 2015; Rodrigues *et al.*, 2016) and that each is thus necessary to predict evolutionary outcomes (Walkiewicz *et al.*, 2012).

Perhaps more importantly, Heinrich *et al.* (1991) and Schuster *et al.* (2008) have pointed out that these modelling frameworks assume a constant value for either or both concentrations of the first substrate and of the final product (Orth *et al.*, 2010), whereas Evolution should instead maximize the amount of final products generated. Klipp and Heinrich (1994) found that the aforementioned concentrations indeed have a major influence on optimal rate constants under certain assumptions. Likewise, nutrient uptake is most often not considered explicitly in existing models of enzyme evolution, while it is obviously critical in the competition for resources (Dykhuizen and Dean, 1994).

Nutrient uptake occurs when metabolites move inwards across cell membranes; it may rely on membrane permeability only (passive diffusion) or involve channels and carrier proteins, be they transporters or cotransporters (Stein, 1986a). Here we build a model that explicitly includes passive (PD hereafter) or facilitated diffusion (FD) followed by an unbranched metabolic pathway to study how resource availability coupled to transport modulates the evolution of enzymes along the pathway. In ecological scenarios where individuals compete for resources, Natural Selection should favour genotypes that maximize the net intake of molecules and their transformation, which are linked under both PD and FD.

Based on this premise, we confirm that the evolution of enzyme kinetic parameters *k_f_* and *k*_cat_ takes place on cliff-like fitness landscapes where a fitness plateau covers a wide part of the relevant parameter space. Kinetic parameters have co-dependent but distinct evolutionary dynamics – and thus distinct sensitivities to certain parameters of the model such that the shape of the plateau can be modulated by changing parameters of the model within realistic ranges. We show that this fitness landscape depends on features of transporters that initiate a metabolic pathway, along with parameters that vary among enzymes within a pathway, like the tolerance to high concentrations of intermediate metabolites or the reversibility of reactions.

We further demonstrate, using a simple population genetics model, that the evolutionarily expected features of an enzyme should be predictable, even though enzymes evolve near-neutrally on the fitness plateau. This is because the model includes slightly biased mutations that tend to produce a majority of less efficient enzymes. We thus postulate that the wide variety of enzyme features reported might be explained in a large part by differences in the shape of their fitness landscapes. While testing this hypothesis will require extensive information about individual enzymes, we made a small step in this direction, showing that enzymes involved in metabolic pathways with different types of transporters exhibit differences that our model qualitatively predicts.

## 2 Results

### 2.1 Passive diffusion is generally inadequate to sustain cell metabolism

In the version of our model in which intake relies on passive diffusion (PD), the net uptake of a nutrient is a direct outcome of its concentration gradient, and therefore of how efficiently the first enzyme catalyses its transformation inside the cell. Assuming that fitness is proportional to the flux of product of this reaction, we find that the fitness landscape has a cliff-like shape with fitness increasing steeply as parameters *k*_cat_ and *k_f_* increase (see Supplementary materials, section Text S1). The precise shape will not be commented in detail here, for it is very similar to landscapes obtained under facilitated diffusion (FD, treated in the rest of this manuscript).

Importantly, our results indicate that PD can only sustain a small part of the metabolism of most living cells given cell permeabilities reported in the literature (Wood *et al.*, 1968; Milo *et al.*, 2010), suggesting that this process may not be a determining factor in the evolution of enzymes along metabolic pathways. Indeed, even extremely efficient enzymes, harbouring values of *k*_cat_ and *k*_cat_/*K*_M_ close to their physical limits, yield low inward fluxes that approach 10^−2^ *mM*.*s*^−1^ when considering a spherical cell with a diameter *D* = 1*μm*. To get a sense of how low these fluxes are, we calculated the maximum cell size they can theoretically sustain. Considering that basal metabolic demands are approximately proportional to the cell volume and using estimates by Lynch and Marinov (2015) for this relationship, we predicted the maximum size enabled by sugar passive diffusion (see 4Materials and Methods). Setting a (conservatively high) medium concentration in glucose [*G*] = 1M yields a theoretical volume ceiling *V_est_* = 0.84*μm*^3^.

Nearly all eukaryotes, and most prokaryotes are *de facto* larger than this threshold (Heim *et al.*, 2017), which might help explain the apparent ubiquity of FD. While this demonstration hinges on numbers for sugar uptake, which may arguably be the task requiring the highest flux, PD may be limiting for many other metabolites (Boer *et al.*, 2010), depending on their permeability and availability in the environment: even for very high amino-acids concentrations that may only be met in multicellular organisms (Schmidt *et al.*, 2016) and assuming the highest observed permeability for such metabolites (Chakrabarti, 1994), these levels are orders of magnitude lower than with FD (see Supplementary material - section Text S1 for PD results).

### 2.2 General shape of the fitness landscape under facilitated diffusion

For most metabolites, FD relies on the specific binding of the substrate to transmembrane carrier proteins (transporters hereafter), followed by its translocation to the other side of the membrane (Danielli, 1954; Kotyk, 1967; Stein, 1986b). Our model builds on ter Kuile and Cook (1994)’s approach to model FD, considering the simplifying assumption of symmetric transport. Within this framework, FD operates on the concentration gradient (Bosdriesz *et al.*, 2018) and is susceptible to saturation, represented by constant *K_T_* – similar to *K_M_* in the Michaelis-Menten equation – and an interaction constant *α* (see Methods for details). We assessed how this saturation phenomenon influences the selection pressure acting on forward enzyme kinetic parameters (*k_f_* and *k*_cat_) under various scenarios.

In order to depict a fitness landscape representative of an average enzyme, we first consider a situation where transporters induce a moderately low rate *V_Tm_* and saturate with a relatively high affinity *K_T_* (FIG. 1, notice that affinity increases when *K_T_* decreases). In this situation, the inward flux at steady-state (which, as argued in the introduction, can be considered representative of fitness) forms a plateau when the upstream enzyme in the metabolic pathway has high *k*_cat_ and *k_f_*. This low equilibrium flux elasticity coincides with the saturation theory (Wright, 1934; Kacser and Burns, 1973; Hartl *et al.*, 1985; Dykhuizen *et al.*, 1987; Dean, 1995; Yi and Dean, 2019), especially with its version incorporating facilitated diffusion (ter Kuile and Cook, 1994; Dean, 1995). The flux plateau is delineated by parallel isoclines (solid and interrupted lines in FIG. 1) oriented in the bottom-right direction of the landscape for intermediate values of *k*_cat_ and *k_f_*, such that decreasing *k_f_* by one order of magnitude can be compensated by a similar increase in *k*_cat_. While this mutual dependency holds even for high *k_f_* values as long as *k*_cat_ is not critically low (*i.e.* when *k*_cat_ > 10^−3^), it stops when *k*_cat_ ≥ 10^3^, where increasing *k*_cat_ no longer improves fitness. Besides, the influence of *k*_cat_ and *k_f_* is not strictly equivalent, since the increase in flux is more gradual in response to *k_f_*.

**Figure 1:**
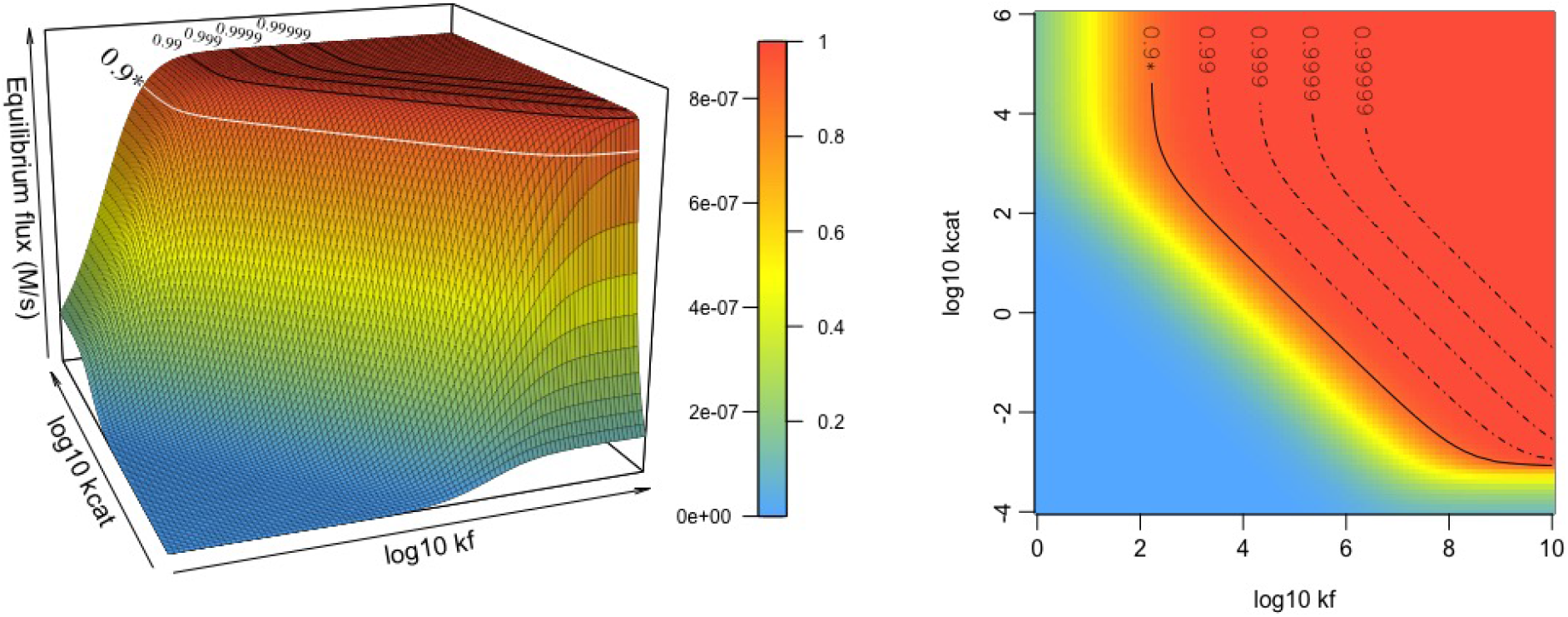
The flux of product following substrate uptake by transporters and conversion by a dedicated enzyme depends on kinetic parameters *k_f_* and *k*_cat_. This landscape is based on a moderately low flux at saturation *V_Tm_* = 1*μM*.*s*^−1^ close to those measured for amino acids and nucleosides in E.*coli* (Zampieri *et al.*, 2019). We also set the transport saturation ratio [*S*_out_]/*K*_T_ to 10 such that the FD process approaches saturation, and relatively high transporter affinity *K*_T_ = 50*μM*, also in line with estimates for nucleosides (Griffith and Jarvis, 1996; Xie *et al.*, 2004). Other parameter values include *k_r_* = 10^3^*s*^−1^ and [*E_tot_*] = 1*mM*. The color gradient indicates the absolute and normalized (with a maximum flux of 1) values of equilibrium flux.

Furthermore, and contrary to the textbook picture whereby most biological reactions are not limited by diffusion at all (Bar-Even *et al.*, 2011; Sweetlove and Fernie, 2018), increasing an enzyme’s association rate *k_f_* – be it through its diffusivity or its binding rate – may still enhance the equilibrium flux when diffusion is substantially faster than catalysis.

### 2.3 Properties of facilitated diffusion modulate the landscape

To explore the effect of FD kinetics on the evolution of enzymes in the metabolic pathway, we studied the influence of *K_T_* – the affinity of the transporter for the substrate – and *V_Tm_* – the maximum transport rate – still assuming that the substrate is close to saturation ([*S*_out_]/*K_T_* = 10). We find that increasing the transport flux *V_Tm_* exerts a positive selection pressure on kinetic parameters for the upstream enzyme (*i.e.* for increasing *k*_cat_ and *k_f_*). The plateau is shifted accordingly (see FIG. 2-A), towards the top-right corner of the landscape, at a distance that corresponds to the magnitude of the change in *V_Tm_*. Increasing the affinity of the transporter (*i.e.* decreasing *K_T_*), however, selects for higher *k_f_* (the isoclines are displaced to the right and the fold change is similar to that of *K_T_*) but has no other visible influence on *k*_cat_ than increasing its codependency with *k_f_*, a result that holds regardless of the flux at saturation *V_Tm_* (notice that we only considered high *V_Tm_*s, larger than in the average case, because these cases are more likely to be under directional selection).

**Figure 2:**
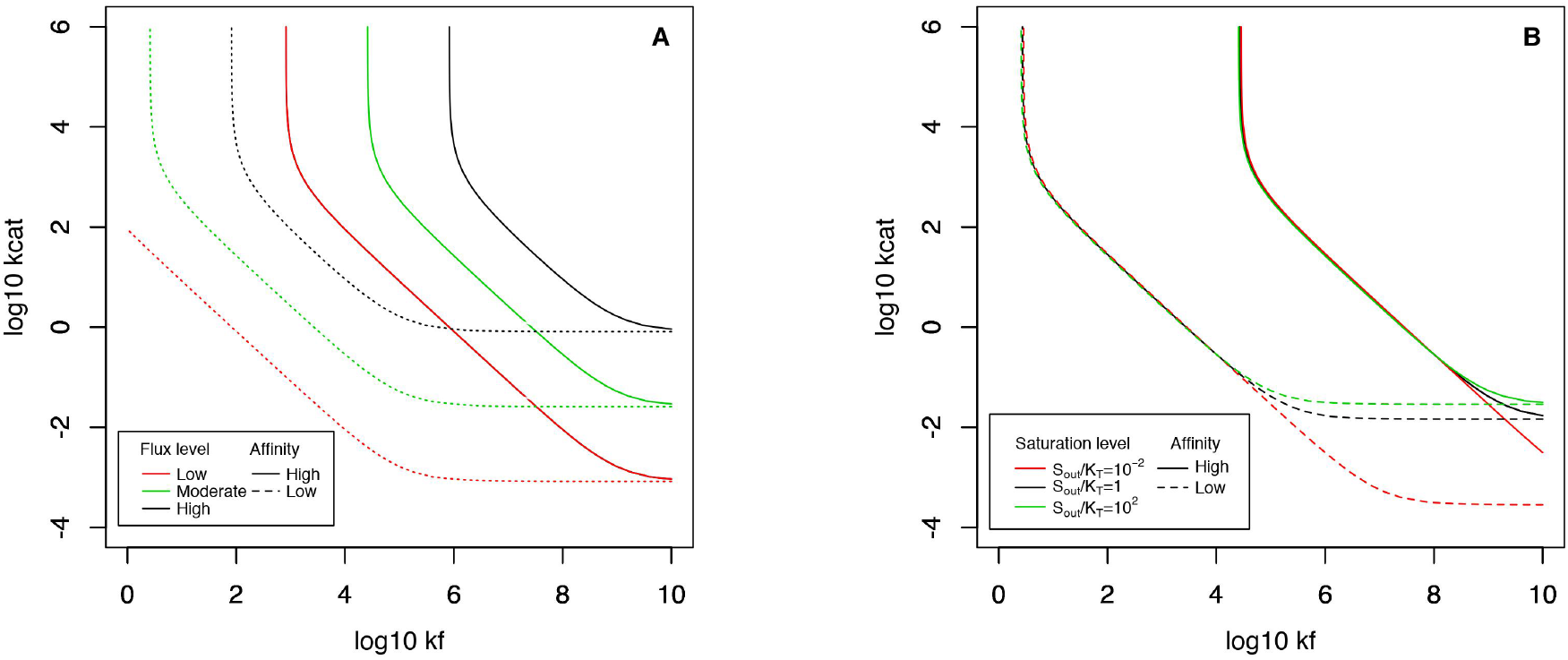
Features of a transporter have an impact on the flux landscape for upstream enzymes, as shown by the 0.9 isoclines – above which the relative flux is > 90% – that delineate the fitness plateau for each set of parameter. A: low (*K_T_* = 0.1*M*) and high (10*μM*) transporter affinities are considered, in combination with low (*V_Tm_* = 10^−6^*M*), moderate (10^−4.5^*M*) or high maximum flux (10^−3^*M*). Increasing *K_T_* extends the plateau only towards the left part of the landscape, allowing enzymes with lower *k_f_* on the plateau, whereas decreasing *V_Tm_* extends the plateau in both directions. B: the shape of the fitness plateau is however little dependent on the saturation of the transporter, for a transporter with moderate flux (*V_Tm_* = 10^−4.5^*M*.*s*^−1^; the effect is identical for higher *V_Tm_*, see SM Fig. S2). Other parameter values: *k_r_* = 1000/*s*, [*E_tot_*] = 1*mM* and [*S_env_*] = 10 × *KT*.

This specific effect on the affinity of the upstream enzyme is likely due to a competition between the transporter – which can transport the substrate in both directions – and the enzyme, which harvests the substrate at a rate that depends on the dissociation constant *K_D_* = *k_r_*/*k_f_*. It should be noted that nutrients under lower demands – *e.g.* amino acids – are generally less concentrated in the environment, often coinciding with a higher affinity of their transporter. Therefore, the possible combinations of flux and affinity likely occupy a restricted space of possibilities where flux and affinity are negatively linked (Gudelj *et al.*, 2010; Bosdriesz *et al.*, 2018), which as can be seen in SM Figs. S3-A,E,I results in landscapes that mainly differ by the minimum value of *k*_cat_ on the plateau. In FIG. 2-A, we have considered ranges of empirical estimates for sugars (high flux with low to moderate affinity) (Stein, 1986b; Maier *et al.*, 2002), nucleosides (Griffith and Jarvis, 1996) and amino acids (Stein, 1986b; Zampieri *et al.*, 2019) (weak to moderate flux with moderate to high affinity), which indeed mainly correspond to these combinations.

So far we have considered transporters saturated by high external substrate concentrations. Relaxing this assumption has little impact on the fitness landscape, except that very low values of *k*_cat_ (lower than 10^−2^ in FIG. 2-B) can only sustain the low influx of transporters far from saturation, but fail to keep up with higher influxes in richer environments.

### 2.4 Enzymes differ among metabolic pathways

We then superimposed empirical estimates of kinetic parameters over our theoretical fitness landscapes, after substituting parameter *k_f_* for its usual empirical counterpart, *k*_cat_/*K*_M_. Because *k_cat_*/*K*_M_ = *k*_*f*_ *k*_cat_/(*k_r_* + *k_cat_*), this approximation only holds when *k*_cat_ ≫ *k_r_*. Representing the fitness landscape in this parameter space sets an inaccessible area in the bottomright part of the landscapes where *k_f_* would exceed the diffusion limit (grey area on FIG. 3). For purposes of inclusiveness, we used *k_r_* = 10^2^*s*^−1^ by default – noting that this limit would be displaced upwards for larger *k_r_* (and downwards otherwise).

**Figure 3:**
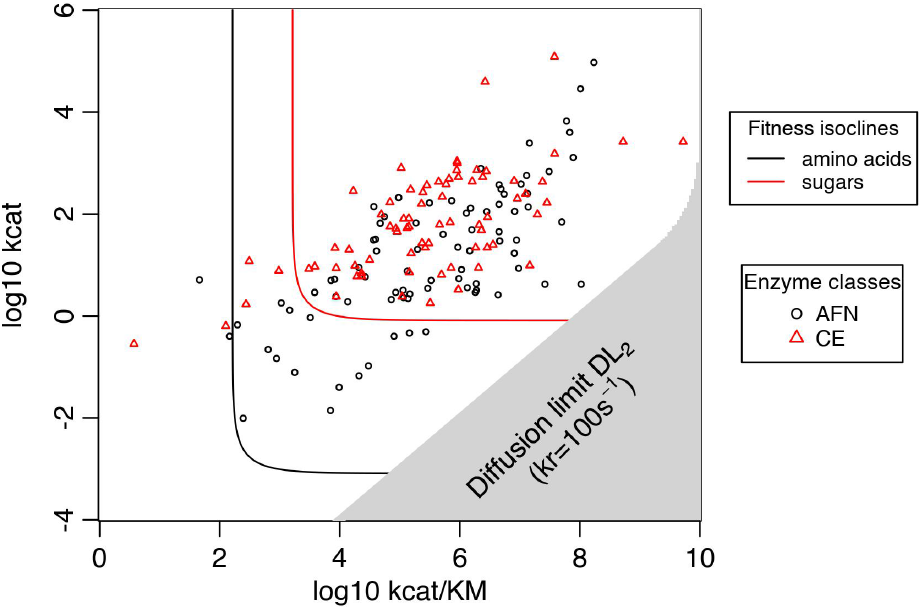

*in vitro* experimental estimates of kinetic parameters *k*_cat_ and *k*_cat_/*K*_M_ exhibit different distributions for enzymes involved in different categories of pathways – as identified by Bar-Even *et al.* (2011) – namely (AFN): amino acids, fatty acids, and nucleotides and (CE): carbohydrates and energy. Corresponding fitness landscapes – differing by transporter features – are superimposed, with the parameter space narrowed down due to the diffusion limit (grey area, set for *k_r_* = 10^2^*s*^−1^). The isoclines shown correspond to parameter values typical of sugar transporters (*K_T_* = 5*mM*, *V_Tm_* = 1*mM*.*s*^−1^, in red) (Maier *et al.*, 2002) or amino acids transporters (same as in FIG. 1, in black).

We otherwise used sets of parameters that correspond to typical features of sugar and amino acid/nucleoside transporters to obtain FIG. 3. Because we have previously shown that changing the affinity or maximum flux of transporters may move the fitness plateau, our model predicts that enzymes involved in the corresponding pathways (*e.g.* of sugars and amino acids) should have their own specific distributions. We see that enzymes involved in the central carbohydrate metabolism as categorized by Bar-Even *et al.* (2011) have on average higher *k*_cat_ and *K*_M_ than those metabolising amino-acids and nucleotides. Our superimposition with the predicted fitness plateaus in FIG. 3 suggests that there may indeed be an explainable difference between enzymes contributing to carbohydrate processing (in red) and to that of other primary metabolites (in black, *e.g.* amino acids). We acknowledge that this result implicitly suggests that enzymes within a pathway have evolved on a common fitness landscape, spreading neutrally onto the fitness plateau. This is by no means our interpretation, as this subset of the full dataset includes enzymes that differ in many other ways that, as we will see, make each enzyme evolve on its own fitness landscape and thereby potentially explain a large part of this observed variance.

### 2.5 Downstream enzymes also evolve on cliff-like fitness landscapes

One of the factors that makes enzymes different along a pathway is their position, such that the fitness landscape in FIG.1 may only hold for the most upstream enzyme in a pathway. Indeed, because the flux of the first product in a pathway increases with the substrate gradient across the cell membrane, the upstream enzyme of a given metabolic pathway is selected for efficiency as described above. In contrast, this selection pressure does not apply directly downstream; at steady-state, even inefficient enzymes can in principle process newly formed substrate molecules at an elevated rate, assuming that the concentration of the substrate is allowed to reach any steady-state value. This is an obviously unreasonable assumption, since a part of this standing substrate should be lost by outward diffusion or degradation (Jones *et al.*, 2015; Bosdriesz *et al.*, 2018). The loss of fitness may therefore result from the loss of metabolites in a way that can be modelled by a constant degradation rate *η_d_* (Chou *et al.*, 2014) (assuming that the external environment is infinite, the degradation term can as well represent an efflux). Highly concentrated metabolites may also be involved in widespread non-specific (Keller *et al.*, 2015) or promiscuous interactions (Khersonsky and Tawfik, 2010; Schäuble *et al.*, 2013; Peracchi, 2018) that may interfere with other cellular processes; this is well captured by the linear cost as non-specific interactions should follow Michaelis-Menten kinetics albeit with much lower affinities, hence following an approximately linear relationship up to very high cellular concentrations (see Materials and Methods for more details). However for some reactions the accumulation of metabolites may result in the production of toxic compounds (Lilja and Johnson, 2017; Niehaus and Hillmann, 2020), hence triggering toxicity best modelled as a non-linear fitness cost (Clark, 1991; Wright and Rausher, 2010).

We first consider a “perfect”, highly concentrated upstream enzyme (*k_f_* = 10^10^*M* ^−1^*s*^−1^, *k_cat_* = 10^6^*s*^−1^, *k_r_* = 10^3^*s*^−1^, [*E_tot_*] = 10^−3^*M*) and focus on the second enzyme in the pathway, showing that it evolves on a fitness landscape that has a similar shape than described above, still hitting a plateau (FIG. 4, with the same parameterization as FIG. 1). The degradation rate creates a ceiling for the concentration of the product of the first reaction, such that reducing *η_d_* allows for higher concentrations (see SM Fig. S4) and makes the flux tolerant to second enzymes with lower *k_f_* s, whereas selection on *k*_cat_ is barely impacted by this parameter. The plateau is therefore extended to the left when high product concentrations are enabled at low *η_d_* (see FIG. 4-B). The shape of the plateau is little impacted by changes in the efficiency of the first enzyme, especially when it stands on the plateau. These results are almost independent of the transporter initiating the pathway (see SM Fig. S6 for the case of moderate affinity, high flux transporters).

**Figure 4:**
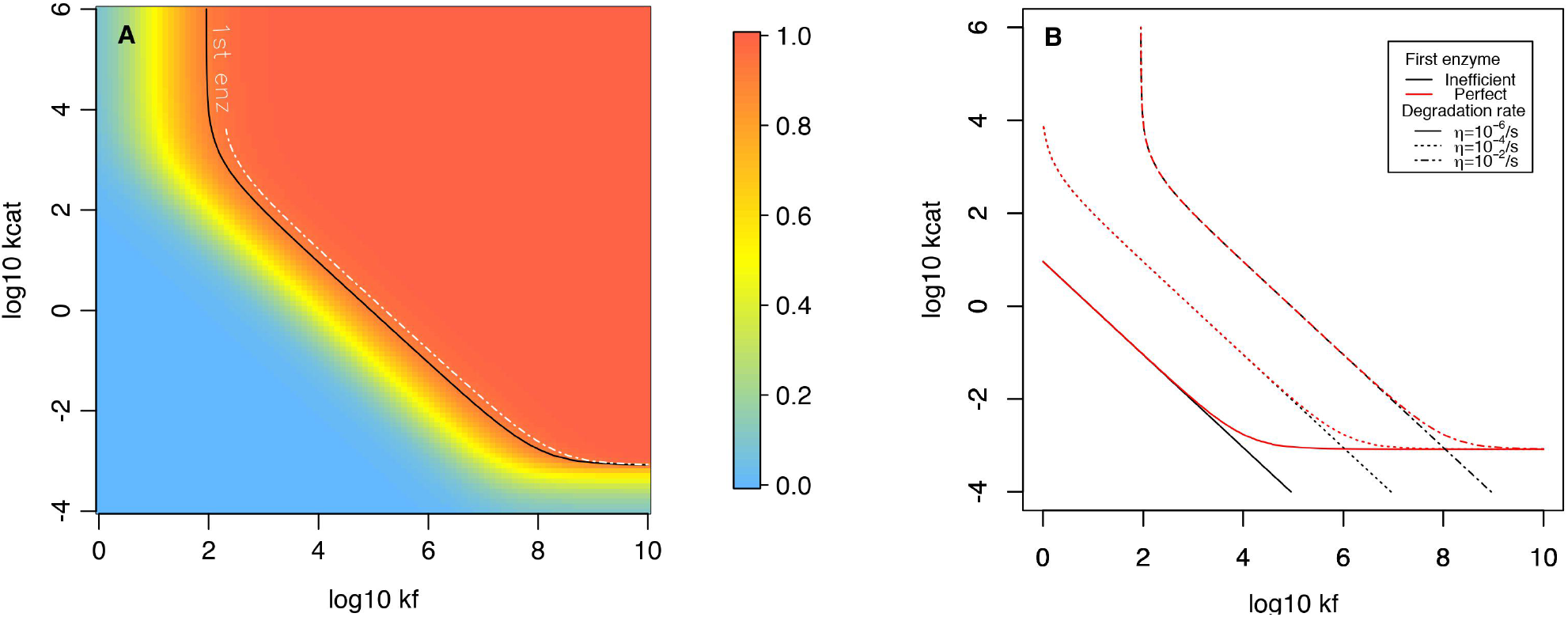

Downstream enzymes exhibit similar fitness landscapes as those upstream, with a dependency to degradation parameter *η_d_*. A: a high degradation rate (*η_d_* = 10^−2^/*s*) results in a fitness plateau for the second enzyme very similar to that of the first enzyme; in the case presented the first enzyme is considered “perfect” in order to draw the fitness landscape of the second enzyme (*k_f_* = 10^10^ *M*^−1^.*s*^−1^, *k_cat_* = 10^6^*s*^−1^, *k_r_* = 10^3^*s*^−1^, [*E_tot_*] = 1*mM*). B: decreasing the degradation rate allows less efficient enzymes (with lower *k*_cat_ or *k_f_*) to reach the fitness plateau. Considering the first enzyme to be inefficient (*k_f_* = 10^2^*M* ^−1^.*s*^−1^, *k_cat_* = 10^−2^*s*^−1^, *k_r_* = 10^3^*s*^−1^) instead of perfect marginally changes the fitness landscape by making organisms tolerant to extremely low *k*_cat_. Other parameter values are identical to FIG. 1 (findings are relatively similar for sugar-like transporters, as reported in SM - Fig. S6).

The shape of the negative relationship between metabolite concentration and fitness can be important (Figs S7-S9 in SM), as it can make the fitness landscape of an enzyme dependent of the overall flux of the metabolic pathway, and therefore on other enzymes in the pathway. Indeed, low general fluxes (as modelled by an inefficient first enzyme in Figs. S7-S8) make the metabolite concentration below its toxicity threshold, therefore making organisms tolerant to enzymes with lower *k_f_* and *k*_cat_. Taken together, these results show that the precise epistatic relation-relationship between enzymes in a pathway will depend on the exact cost function applied, with a linear cost generating epistasis for *k*_cat_ only and a non-linear cost possibly impacting both *k_f_* and *k*_cat_.

### 2.6 The reversibility of reactions also matters

Reversibility is an intrinsic feature of chemical reactions that cannot be directly overcome by Evolution (Haldane, 1930; Cornish-Bowden, 1979). A highly reversible reaction corresponds to a large intrinsic equilibrium constant *K_eq_* = [*S*]_*eq*_/[*P*]_*eq*_ (Klipp and Heinrich, 1994), and results in higher backward than forward rates in the following chemical equation:

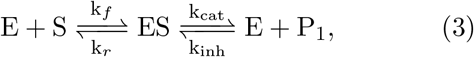

where *k_inh_* represents the rate at which enzyme and product combine back. Such a (reversible) reaction could in principle influence the selective pressure acting on the following enzyme in the pathway, for both enzymes compete to process the same metabolite *P*_1_. We thus quantified how reversibility affects the evolution of an enzyme downstream (SM Figs S10 and S11).

The equilibrium constant *K_eq_* has a similar (non-linear) impact on the fitness landscape of the second enzyme to that of the degradation rate, with a highly reversible upstream enzyme exerting a selection pressure downstream towards an increase of kinetic parameters (SM Fig. S10-A). Indeed, increasing *K_eq_* moves the fitness plateau toward the upper-right corner in the (*k_f_*, *k*_cat_) parameter space, hence selecting for more efficient downstream enzymes. The effect appears linear, except for very low values of *K_eq_* where metabolite accumulation exerts a dominant role in shaping the fitness landscape (through the degradation rate *η_d_*, set to a low residual value). Therefore, the reversibility of the upstream reaction appears like a critical parameter for the evolution of an enzyme.

### 2.7 Evolutionary dynamics of enzyme kinetic parameters

How much variance in evolutionary outcomes these differences in fitness landscapes may explain is contingent on the interplay between selection, mutation and drift. Small differences in an isocline position should indeed be of little importance if populations perform random walks on the fitness plateau, for instance. To approach how populations evolve on our mathematically derived fitness landscapes, we built a simple population genetics model in which absolute fitness is directly proportional to the flux arising from the first enzyme at steady-state – which itself equals the net inward flux of nutrients. Two different levels of metabolic demands were considered, corresponding to parameter values of amino acids/nucleosides and sugar transporters (panels (A) and (I) in SM Fig. S3). In this instance of the model, only *k*_cat_ and *k_f_* were susceptible to evolve through mutations. Mutational effects on log_10_ *k_cat_* and log_10_ *k_f_* were drawn from independent normal distributions with mean *b* ≤ 0, and the absolute value of *b* setting the intensity of a mutational bias towards less efficient parameter values, as has been widely documented in many contexts (Eyre-Walker and Keightley, 2007; Serohijos *et al.*, 2012; Heckmann *et al.*, 2018). The standard deviation of the distribution of mutational effects equals 0.3 such that most mutations explore the neighbouring parameter space, seldom changing a parameter by more than one order of magnitude (one log_10_ unit) in compliance with empirical estimates (Carlin *et al.*, 2016). Since the relation between kinetic parameters may be constrained – *e.g.* due to shared properties of the energy profile of a reaction – we tested the influence of negative and positive relationships using bivariate normal distributions, with three different values of *ρ* (see Materials and Methods for details).

In the absence of mutational bias (*b* = 0), simulated enzymes spread over the fitness plateau, as expected (Fig. S16-A for low flux, Fig. S17A otherwise). The onset of the plateau depends on the strength of drift and hence derive from the effective population size *N_e_*, following the classical expectation that selection becomes inefficient when *N_e_* × *s* < 1 (Wright, 1931; Kimura, 1968). Introducing a mutational bias that makes enzyme kinetics less efficient on average has a strong effect on both *k*_cat_ and *k_f_*, preventing simulated enzymes from improving far above the drift barrier (FIG. 5-A for low flux, FIG. 5-B otherwise). Even weak biases (*b* = −0.1) lead to enzymes evolving in the vicinity of the isocline where *N_e_* × *s* ≈ 1. Increasing the strength of this bias to 0.2 only slightly decreases the population variance around this expectation. Finally, mutational correlations do not impact much the distribution of evolutionary outcomes (SM Fig. S18).

**Figure 5:**
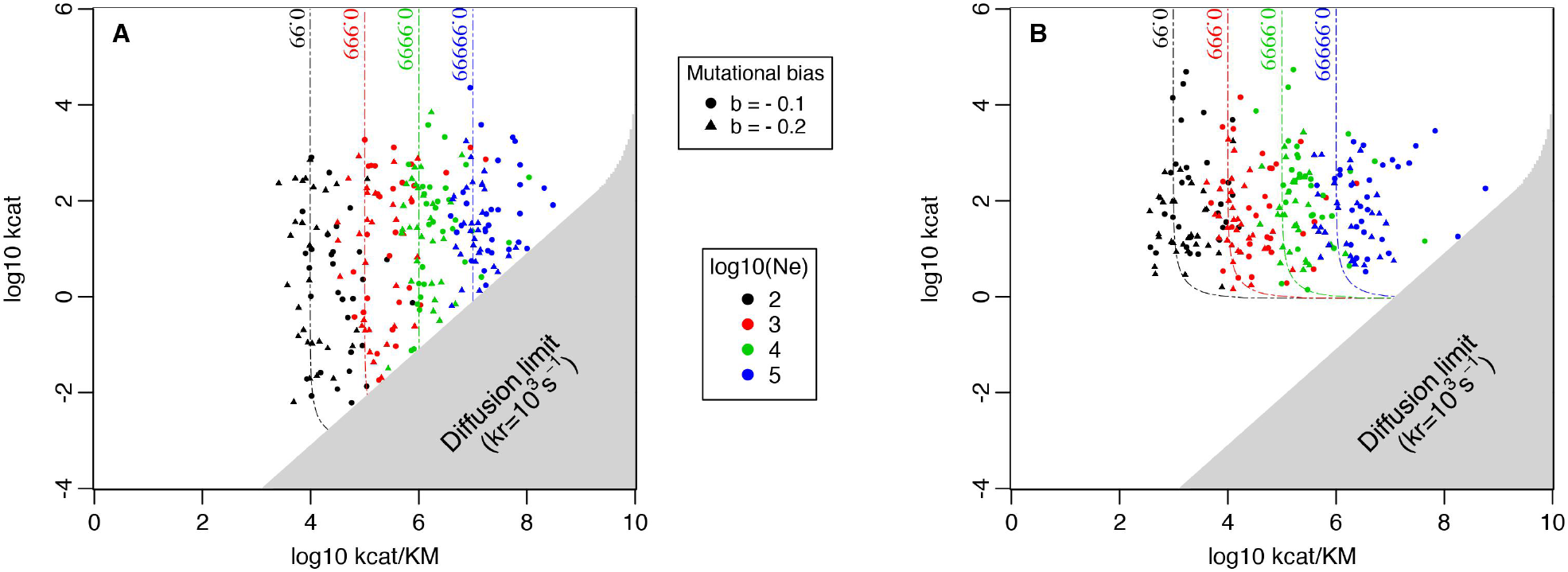
Population genetic simulations predict that enzymes should reach a predictable set of features when mutation biases towards lower efficiencies are considered (see SM fig. S17 for the case of an absence of bias). Indeed, the mutation selection drift equilibrium establishes close to an isocline indicative of effective selection that depends on the effective population size *N_e_*. The cases considered here are that of a transporter with a low flux at saturation and high affinity (A; *V_Tm_* = 1*μMs*^−1^ and *K_T_* = 10*μM*) and one with a high flux at saturation but low affinity (B; *V_Tm_* = 1*mMs*^−1^ and *K_T_* = 100*mM*) with effective population sizes ranging from 10^2^ to 10^5^ (different colors) and two strengths of the mutational bias (the absence of mutational bias was also considered, see SM). Each of 30 independent simulations for each scenario is represented a dot in the “empirical” parameter space (*k*_cat_, *k*_cat_/*K_M_*), but only *k*_cat_ and *k_f_* were susceptible to evolve. *k_r_* is set to 10^3^*s*^−1^ such that the grey part of the parameter space is inaccessible to enzymes that would otherwise exceed the diffusion limit.

Our results suggest a strong effect of the effective population size on enzyme evolution, such that species with *N_e_* above 10^5^ (Bobay and Ochman, 2018, most unicellular organisms) should express extremely efficient enzymes. This appears to not be the case, as for instance Eukaryotes and Prokaryotes display similar enzymes despite large differences in effective population sizes (Bar-Even *et al.*, 2011). As we will later discuss, this conundrum might resolve when considering the smaller size of organisms forming large populations, making them more sensitive to noise in gene expression and favouring higher concentrations. Notwithstanding this issue, the prediction of enzymes evolving a predictable set of kinetic parameters strongly suggests that a large part of the broad variance in enzyme features is due to differences in the selective context experienced by each, thereupon requiring further investigation on the dependency of the position of the fitness plateau to parameters of our model.

### 2.8 The joint evolution of enzyme concentrations and kinetic parameters

Hitherto, we have considered enzymes to be highly concentrated, an assumption that we now relax since it is an important component of the presumed kinetic activity (Koshland, 2002). Predictably, increasing the concentration of the first or second enzyme in a pathway releases the selection on their kinetic parameters (Noor *et al.*, 2016), producing larger fitness plateaus as an enzyme concentration increases (see SM - Figs S12-B and S13-B for this influence in different contexts). Due to the compensatory effects between concentration and activity, we anticipate that the joint evolutionary dynamics of the concentration and kinetic parameters should yield a negative correlation between them, as reported by Davidi *et al.* (2016, 2018).

Despite their common role on reaction efficiency, enzyme concentration expectedly responds to very different selection pressures than kinetic parameters, as increased gene expression levels come with costs (Wagner, 2005; Lang *et al.*, 2009; Scott *et al.*, 2010; Noor *et al.*, 2016; Kafri *et al.*, 2016). Indeed, producing extra proteins requires both energy and matter (Novick and Weiner, 1957; Stoebel *et al.*, 2008; Wagner, 2005; Lynch and Marinov, 2015) and may impede the efficiency of physical processes that rely on an optimal intermediate content (Dong *et al.*, 1995; Dill *et al.*, 2011; Andrews, 2020). We designed a new instance of our population genetics model to study the tangled evolution of kinetic constants and enzyme concentration, introducing two of these costs: (1) the cost of producing proteins *c*, considered to be proportional to concentration (Wagner, 2005; Chou *et al.*, 2014; Lynch and Marinov, 2015), and (2) the exponential cost of an increase in macromolecular crowding, which hinders diffusion and thus slows down reactions (Dill *et al.*, 2011; Schavemaker *et al.*, 2018; Andrews, 2020) (see SM Fig. S15 for the resulting fitness landscapes of enzyme concentration).

The two types of costs result in a different shape of the fitness landscape, with the noticeable difference that evolutionarily expected concentration depends on *N_e_* when the cost of production is considered (SM - Fig. S19) but not with crowding effects (SM - Fig. S20). With a combination of the two costs, enzyme concentrations decrease with *N_e_* and production costs, resulting in the evolution of higher kinetic constants (FIG. 6). This is because at higher effective sizes, direct costs of protein production are large enough to incur effective selection for lower protein expression. This is no longer the case when *N_e_* decreases, such that the major force driving the optimization of enzyme concentration becomes that opposing macromolecular crowding, which is less sensitive to *N_e_* (as shown in Fig. S19 in SM). The balance between these two selective forces, and the dependency to *N_e_*, obviously depend on the relative importance of these costs (SM - Fig. S20), itself depending on many parameters (protein length, molecular weight, etc.) that should only make enzymes marginally different within a given species (when their activity evolves on similar fitness landscapes).

**Figure 6:**
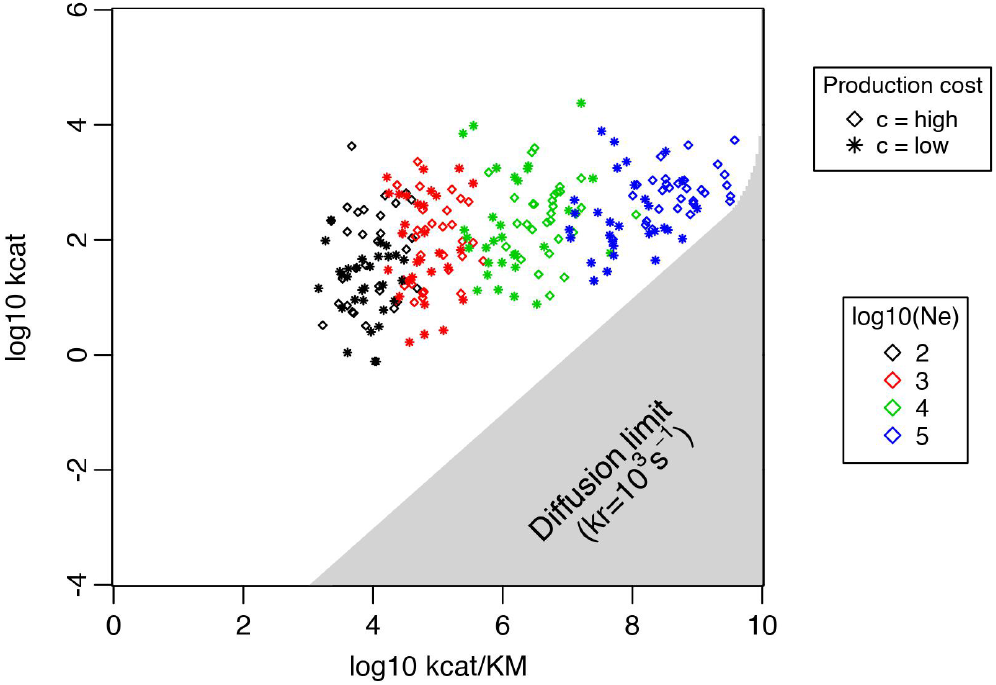
Simulations of the joint evolution of enzyme concentration and kinetic parameters, with a twofold cost of enzyme overexpression (the direct metabolic cost and the indirect cost of cell packing). The case considered here is that of a transporter with a high flux at saturation and low affinity (*V_Tm_* = 1*mMs*^−1^ and *K_T_* = 1*mM*) under a high mutational bias on kinetic constants (*b* = −0.2). Two different costs of protein production *c* are considered along with four effective population sizes ranging from 10^2^ to 10^5^. We ran 30 independent simulations for each scenario, each represented by a dot in the “empirical” parameter space as described in fig. 5.

## 3 Discussion

Most enzymes have been considered to be only moderately efficient (Bar-Even *et al.*, 2011), if not sloppy (Bar-Even *et al.*, 2015). This claim was put into perspective by Newton *et al.* (2018) who argued that the link between fitness and enzyme efficiencies is complex and may be partly enzyme dependent, such that all enzymes may not evolve on a common fitness landscape. Through this work, we have developed a model where enzyme efficiencies are mechanistically linked to fitness through the impact of nutrient gradients on the production of metabolites. Our results emphasize that an enzyme’s fitness landscape – and the resulting mutation-selection-drift balance – may indeed be largely context dependent, possibly explaining a large part of the extreme observed variance in enzyme features.

At first sight, all enzymes evolve on fitness landscapes that have the same general shape, with a fitness plateau surrounded by a steep slope. While this shape is usual in models of enzyme evolution (Hartl *et al.*, 1985; Kaltenbach and Tokuriki, 2014; Yi and Dean, 2019), in our model the landscape is drawn in the parameter space formed by the two forward kinetic parameters *k*_cat_ and *k_f_*, instead of a composite “efficiency” whose relevance is questionable (Eisenthal *et al.*, 2007; Koshland, 2002). Our model allows to predict the precise position of the fitness plateau in various contexts, showing that model parameters may have a selective impact on *k_f_*, *k*_cat_, or both, thereby confirming the relevance of considering their distinct evolutionary dynamics.

We have shown that the exact position of the plateau is important through a population genetics model including mutational biases that produce less efficient enzymes at a slightly higher frequency. Despite their small effect, these biases are sufficient to have a significant impact on the evolutionary dynamics occurring on the fitness plateau, preventing enzymes to explore the parameter space far away from an isocline whose precise value can be predicted. Because the mutation-selection-drift balance occupies a narrow part of the landscape, this makes the evolution of an enzyme, in principle, highly predictable. Likewise, we anticipate that differences between enzymes should largely be explained by differences in the shapes of their individual fitness landscapes.

Overall, the selective pressure acting on an enzyme results from an interplay between several biochemical factors. We have effectively found that the shape of the fitness landscape is first governed by features of the transporter initiating a pathway, especially the maximum flux they can sustain. Using parameters that correspond to empirical estimates for sugars and amino acids/nucleosides, we have found that enzymes contributing to subsequent metabolic pathways should be different, with those in the “sugars” pathway being selected for faster kinetics.

While sharing a common transporter, enzymes along a pathway are also subject to a variety of local chemical contexts, making each evolve on its own unique fitness landscape. This could explain, at least in part, the large within-pathway variance of enzyme kinetic parameters. Physical constraints may for instance act differentially on different enzymes, as exemplified by the intrinsic reversibility of a reaction that fuels the selective pressure towards higher efficiency in downstream enzymes. This may contribute to explain the high efficiency of a few enzymes like TIM (Williamson *et al.*, 1967; Davidi *et al.*, 2018).

One way to compensate for low kinetic constants is to enhance the level of expression of an enzyme, as confirmed by our model – concentration indeed has a strong influence on the fitness landscape of *k_f_* and *k*_cat_. Nonetheless, concentration and kinetic parameters face very distinct selection regimes: while the latter are both under directional selection, vanishing at high efficiencies, concentration is under stabilizing selection – owing to a combination between its positive impact on the flux and the adverse costs to high expression – as already pinpointed by Chou *et al.* (2014). Their joint evolution is complex because the position of the concentration optimum depends on an enzyme’s kinetic constants, whose fitness landscape itself depends on concentration. This results in a slightly increased variance in enzyme efficiencies compared to simulations with fixed concentrations, along with a complex relationship with genetic drift, because small populations tend to tolerate higher enzyme concentrations and, therefore, evolve less efficient enzymes.

It should be noted that our model does not consider another selection pressure on enzyme concentrations that arises from noise in gene expression, as argued by Wang and Zhang (2011). Indeed, low expression results in detrimental noise that should be avoided by pushing enzyme concentrations towards higher values in small organisms like Prokaryotes (see SM section Text S6 for an estimate of this effect). This could result in a different relationship between *N_e_* and enzyme efficiencies than considered here, possibly explaining the confusing observation that species with larger populations (and smaller sizes) do not express markedly more efficient catalysts. Furthermore, an enzyme’s effective concentration can also increase through compartmentalization (Ovádi and Saks, 2004; Diekmann and Pereira-Leal, 2013; Cornejo *et al.*, 2014) and substrate channeling (Welch and Easterby, 1994; Huang *et al.*, 2001; Sweetlove and Fernie, 2018), within the limits imposed by noise, and modify the selective pressure acting on kinetic parameters.

This illustrates that rather than making precise predictions, our study aims at making the strong claim that selection acting on enzyme features is important for their diversity, possibly largely overcoming the diversity arising from neutral processes. If this is indeed the case, trends in enzyme evolution can be predicted – as it was shown empirically in the context of antibiotic resistance (Walkiewicz *et al.*, 2012) – and further improvements of this model should allow to predict the expected features of individual enzymes. Such improvements are made easier by the use of a mechanistic framework, where fitness arises as enzymatic efficiency helps ingesting nutrients and win the competition for resources. This framework could even be enriched by other dimensions relevant to the genotype-phenotype-fitness map (Bershtein *et al.*, 2017; Echave, 2019; Kinsler *et al.*, 2020).

Unfortunately, mechanistic does not mean free of a definition of fitness, as we have here assumed that the latter is proportional to metabolic flux, hence considering each flux in isolation. Fitness instead results from a wide range of metabolic pathways that combine together and should all be competitive to certain degrees. How global epistasis builds up (Weinreich *et al.*, 2013; Otwinowski *et al.*, 2018; Reddy and Desai, 2020), and genetic drift acts in this context, is far from obvious (Iwasa *et al.*, 2004; Weinreich and Chao, 2005; Weissman *et al.*, 2009). But this should not impact much how enzymes evolve in old, overall efficient pathways, as any impediment in efficiency should have a relatively independent effect on fitness in this context, as captured by our model. Understanding these complex interactions between pathways would nevertheless be crucial to understand how metabolic pathways arose and improved, likely from a highly inefficient state during early steps in the evolution of life on Earth (Kacser and Beeby, 1984; Schmidt *et al.*, 2003; Heckmann *et al.*, 2018).

## 4 Materials and Methods

### 4.1 Quantifying the maximum size for cells using passive diffusion

If a cell is to be viable, it has, at least, to uptake enough glucose to compensate for basal metabolism – metabolism that allows to maintain the same cell size for non-actively growing cells (Lynch and Marinov, 2015) – leading to the following equation: *ϕ_PD_* = *C_M_*, with *ϕ_PD_* the uptake through passive diffusion and *C_M_* the basal metabolism demand. To calculate the maximum size a cell can reach using only passive diffusion, we relied on the formula *C_M_* = 0.39*V* ^0.88^(10^9^*ATP/hr*) estimated in (Lynch and Marinov, 2015). We also assumed the cell to be of spherical shape, both concentrations – inside and outside the cell – to be constant with the cellular concentration staying so low that it can be overlooked, meaning that the uptake resulting from passive diffusion can merely be written as 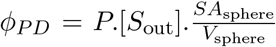, where *SA*_sphere_ and *V*_sphere_ are the surface area and the volume of a sphere, and *P* represents the cell permeability and was measured to 10^−6^*μm*^−1^ (Wood *et al.*, 1968) for glucose. Finally, we considered that each glucose yields 30 ATP molecules (Rich, 2003).

### 4.2 Flux sustained by the first enzyme

When assessing the flux of product made by the first enzyme in a pathway, both (PD) and (FD) result in similar sets of equations; we focus on FD here (see Text S5 - Mathematical appendix in Supplementary material for a comparison with PD). FD typically relies on the specific binding of substrate molecules – located outside the cell – by transmembrane carrier proteins, followed by their translocation into the cytoplasm (Danielli, 1954; Wilbrandt and Rosenberg, 1961; Kotyk, 1967; Bosdriesz *et al.*, 2018). This specific process obeys Michaelis Menten-like kinetics when transport is assumed to be symmetric (Kotyk, 1967), which can be modelled through Briggs-Haldane equations (Briggs and Haldane, 1925; Haldane, 1930; Stein, 1986b):

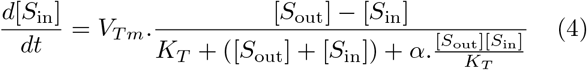

with:

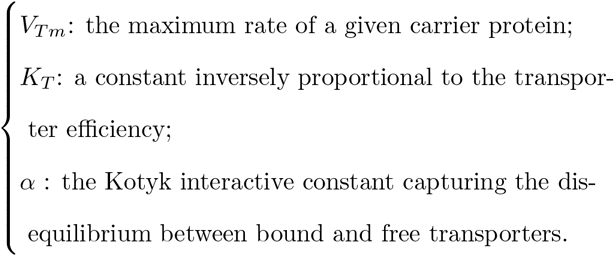

By construction, *α* cannot exceed 1 Kotyk (1967) and is close to this upper limit for sugars (e.g. *α* = 0:91 for glucose (Teusink *et al.*, 1998), so we set *α* = 1 by default in this study, maximizing the effect of interaction).

A model including both FD and irreversible substrate conversion by an enzyme therefore corresponds to the following chemical equation:

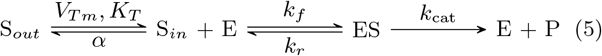

Using analytical tools (see ter Kuile and Cook (1994) and Bosdriesz *et al.* (2018), rederived in Supplementary material - Text S5 Mathematical appendix), the flux can be determined through the following set of equations:

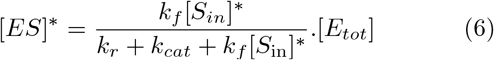

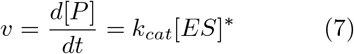

where:

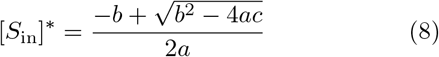

with:

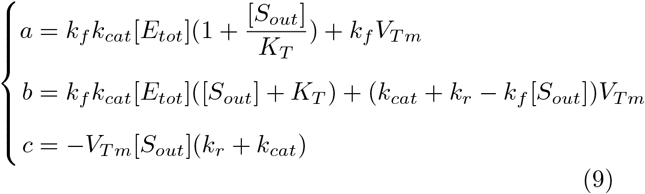

### 4.3 Multiple enzymes model

In order to study the evolution of downstream enzymes, we considered an unbranched metabolic pathway in which the product formed by the first reaction serves as the substrate for a second reaction whose flux determines fitness. Theoretically, as there is nothing prohibiting increase in product concentrations – for it is not considered reversible at this point – any second enzyme should be able to sustain any metabolic demand. We penalized large increases in cellular concentrations through a degradation process of the product of the first reaction, occurring at rate *η_d_* (× this concentration). The chemical reactions occurring after uptake (Michaelis Menten part of Eq.5) are described by the following equations:

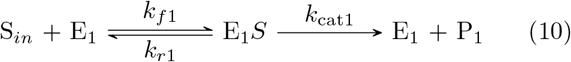

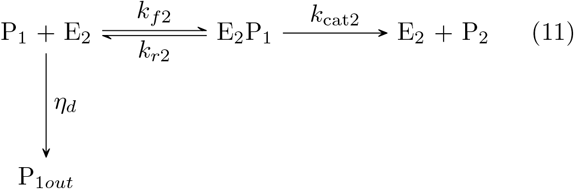

Such a system may reach a steady-state at which the cellular concentrations of the substrate *S_in_* and of the first product *P*_1_ are constant. At this point, the net instantaneous uptake of substrate equals the instantaneous production of *P*_1_ which, in turn, equals the sum of the instantaneous amount of degraded *P*_1_ and the instantaneous production of *P*_2_, according to the principle of mass conservation. It yields the following system of equations:

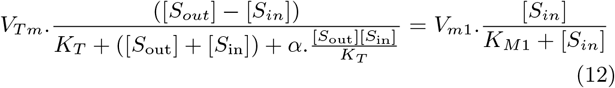

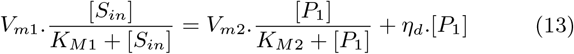

where appear the traditional Michaelis-Menten kinetic parameters for the (i^*eth*^) reaction:

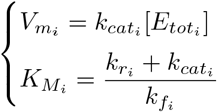

To test the potential influence of toxicity, we defined the absolute fitness as a function of both the flux and a toxicity factor whose influence is represented through a sigmoid function and reflects the non-linearity nature of toxic effects (Clark, 1991; Wright and Rausher, 2010): 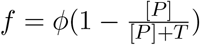

In an independent section, we also introduced reversibility through the modification of equation (10), which becomes:

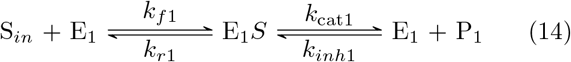

Such a phenomenon is described by the more general form of Haldane equation (Haldane, 1930; Cornish-Bowden, 1979), which changes the contribution of the first reaction 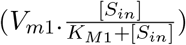 in both equations (12) and (13) to:

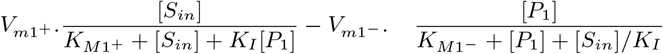

with *V*_*m*1+_ and *K*_*M*1+_ respectively correponding to the previous *V*_*m*1_ and *K*_*M*1_, while:

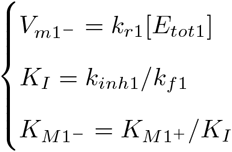

To solve these systems we implemented the Newton method (Atkinson, 1989) aiming to find [*S_in_*]^∗^ and [*P*_1_]^∗^. We ran the algorithm starting from very low values of concentration (both set to 10^−20^*M*) to determine numerically the equilibrium without facing convergence problems. The final flux can then be determined through the “production” part of equation (13), *i.e.* 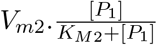.

### 4.4 Validation of the method and computing of the fitness landscapes

To validate the approach, we compared equilibrium results obtained with Raphson-Newton algorithm to those obtained when simulating the process with Euler explicit schemes for a set of (3×3) kinetic values – *k*_cat_ and *k_f_* – encompassing three orders of magnitude (see SM - Section Text 5 for further details).

We then drew fitness landscapes after determining the flux – assumed to be to be linearly related to fitness – achieved for enzyme kinetic parameters *k*_cat_ and *k_f_* varying by 10 orders of magnitude, setting *k_r_* to 10^3^*s*^−1^ – within the range found for several enzymes (Klipp and Heinrich, 1994; Knowles and Albery, 1977) – unless stated otherwise. Except in the section dedicated to the influence of enzyme concentration, we set the enzyme concentration such that [*E_tot_*] = 1*mM*, lying nearby the highest observed values (Albe *et al.*, 1990; Noor *et al.*, 2016). Other parameters are detailed on a case-by-case basis as they may change depending on the goal of each section. To compare with the data and visualize results in the experimenter’s parameter space, we also determined the flux and plotted simulation results using *k*_cat_ and 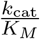, also making them vary by 10 orders of magnitude. We divided the parameter space in 100 log-equidistant values (250 for representations with *k_cat_/K_M_* to obtain a cleaner demarcation for the diffusion limit).

### 4.5 Population genetics model

Evolutionary simulations rely on a Wright-Fisher process including the selective effects of mutations displacing enzymes on mathematically-derived fitness landscapes. Two fitness landscapes were considered: weak flux, high affinity (Fig. S3 A of SM) and high flux, low affinity (Fig. S3 I of SM), both with saturated facilitated diffusion ([*S_out_*] = 10*K_T_*) and the following constant parameters: *k_r_* = 10^3^*s*^−1^ and [*E_tot_*] = 1*mM*. Mutations occur at a rate *μ* = 10^−1^/*N_e_* following reproduction, with an effect sampled in Gaussian distributions with dispersion (*σ* = 0.3). The mean effect of a mutation is given by parameter *b*, hence representing the mutation bias – absent with *b* = 0, low (*b* = −0.1) or high (*b* = −0.2). Kinetic parameters were initiated to the inefficient values of *k_cat_* = 10^−3^*s*^−1^ and *k_f_* = 10^2^*M* ^−1^*s*^−1^ and *k_f_* was limited to values under the diffusion limit – 10^10^*M* ^−1^*s*^−1^ (*k_cat_* was also limited to 10^6^*s*^−1^ when *b* = 0 to avoid physical outliers). To analyse simulation outcomes, we picked out the kinetic and fitness values of the most represented genotype when multiple variants were segregating. 30 simulations were ran for each set of parameters. Finally, we verified that evolutionary steady-states were reached and considered it was the case when at least the average fitnesses (over all simulations) of the last three time-steps were not significantly different one from another (SM Figures S5 and S6).

We also simulated the case where mutations between parameters are correlated. We tested both positive and negative mutational relationships through a bivariate Gaussian distribution whose correlation coefficient were set to *ρ* = −0.8, *ρ* = −0.5, *ρ* = +0.5 (see SM Figure 18 for the results with a moderate flux).

### 4.6 Evolution of enzyme concentrations

Finally, we simulated the joint evolution between kinetic parameters and enzyme concentration, repeating the previous procedure with concentration as an evolvable quantity and the fitness function including deleterious effects of extra gene expression (see SM section Text S5 for the effect of each cost considered independently one from another). Mutations affected either levels of expression or kinetic constants, with those affecting levels of expression (in log values) being drawn from Gaussian distributions with mean 0 and *σ* = 0.2 to comply with existing estimates (Landry *et al.*, 2007; Metzger *et al.*, 2016; Hodgins-Davis *et al.*, 2019). Because sugars are directly involved in energy metabolism, we computed these simulations for the case of a high flux which can more readily be linked to the cost of expression.

As explained above, producing extra proteins is costly, both energetically and because it may increase a cell’s crowding. The cost of protein production is considered to be proportional to the steady-state enzyme concentration, with a slope *c*. Empirical estimates suggest that *c* should be in the range [10^−4^, 10^−3^] (Wagner, 2005; Lynch and Marinov, 2015), such that producing an extra mM of a protein would impede the whole energy budget by one 10000^th^ to one 1000^th^ (we also consider *c* = 10^−5^ and 10^−2^ in the SM). Accordingly, we calculate the absolute fitness *f* = Φ − [*E_tot_*]*c*, where Φ is the flux generated by the enzyme.

The influence of crowding was computed by penalizing *k_f_* through an effective 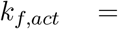 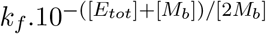, where [*M_b_*] = 2.5.10^−3^*M* represents the basal protein concentration of a viable cell and also constitutes a scaling factor that complies with Andrews (2020) log-linear influence of crowding or glass transition effects described by (Dill *et al.*, 2011).

### 4.7 Data availability

All the enzyme data used in this work to compare fitness landscapes and measured values were recovered from (Bar-Even *et al.*, 2011), and so was the classification of reactions with regards to metabolic groups. Thanks to their authors and publisher, datasets are publicly available at https://pubs.acs.org/doi/10.1021/bi2002289. Apart from that, no new data were generated in support of this research.

## Supporting information

Supplementary Materials for Resource Uptake and The Evolution of Moderately Efficient Enzymes

## 5 Supplementary Material

Supplementary material is available online.

