## Supplementary Materials for Resource Uptake and The Evolution of Moderately Efficient Enzymes for "Resource uptake and the evolution of moderately efficient enzymes"

#### Text S1 Fitness landscapes with passive diffusion

Passive diffusion (PD) efficiency is proportional to the surface-to-volume (SA:V) ratio such that smaller cells are more efficient than larger ones. Here we focus on rather small cells ($D = 1\mu m$), alike Prokaryotes rather than Eukaryotes, whose larger ratio befits this mode of diffusion. Fitness landscapes for upstream enzymes are represented in Figure S1.

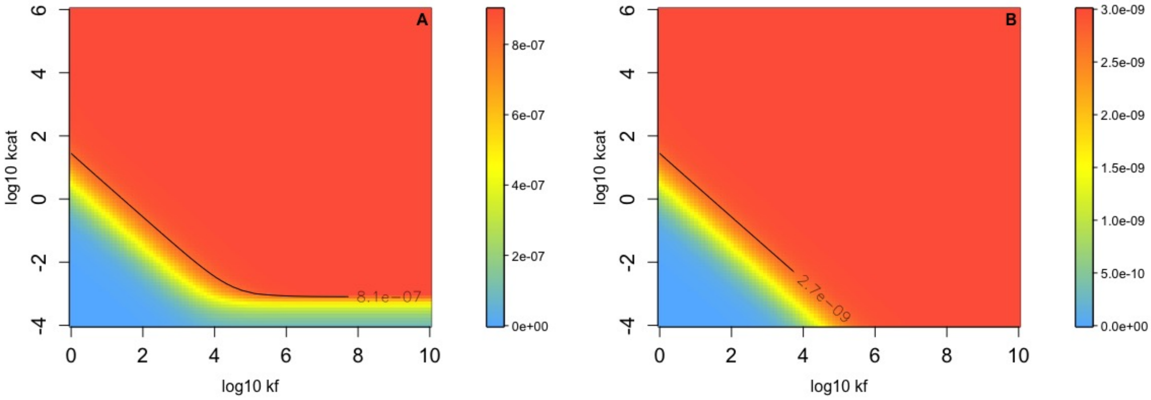


Figure S1: Fitness landscapes for the first enzymes in pathways when uptake results from PD with $P = 10^{-12}m^{-1}$(Wood *et al.*, 1968; Chakrabarti, 1994) in line with values for nucleosides or sugars for instance. The isocline corresponds to 90% of the maximum flux. In panel A, the flux plateau is shown for an abnormally high environment nucleoside concentration ($[S_{out}] = 0.15M$) – that would be more consistent for sugars – which allows to compare the respective landscapes for PD and FD when similar levels are reached, although the situation described for PD is thence far-fetched. The trend – steep slope, large plateau – is even more pronounced than for FD, because concentrations used with FD are low in comparison, which proved to move the plateau towards higher values for both kinetic parameters. In panel B, the flux is represented given the same concentration as that used with FD ($[S_{out}] = 0.5mM$), showing a large plateau whose span is barely demarcated by an area sustaining a very low flux. Furthermore, the flux obtained is more than 2 orders of magnitude lower than for facilitated diffusion, suggesting that PD can only marginally contribute to nutrient uptake even under an extremely favourable scenario.

Comparing with facilitated diffusion (FD, shown in the paper), one can see that for a similar flux (Fig S1-A) the landscape is even flattest with PD, because of the flattening influence of high external concentrations (for a same flux level) that increase nutrient gradients. Anyhow, the substrate concentration yielding such a high level is either very unlikely to be found even within multicellular organisms – for nucleosides/amino acids – or still yielding very low flux relatively to FD – for sugars – as sugar flux are many orders of magnitude higher than nucleosides/amino acids ones (see (A) of Figure S2 for absolute flux with sugar-like transporters). Likewise, setting the concentration to more realistic levels for nucleosides/amino acids does not alter the shape of the landscape while proving PD to be nowhere near the efficiency of FD (illustration (B) on the right of Figure S1). Besides, as

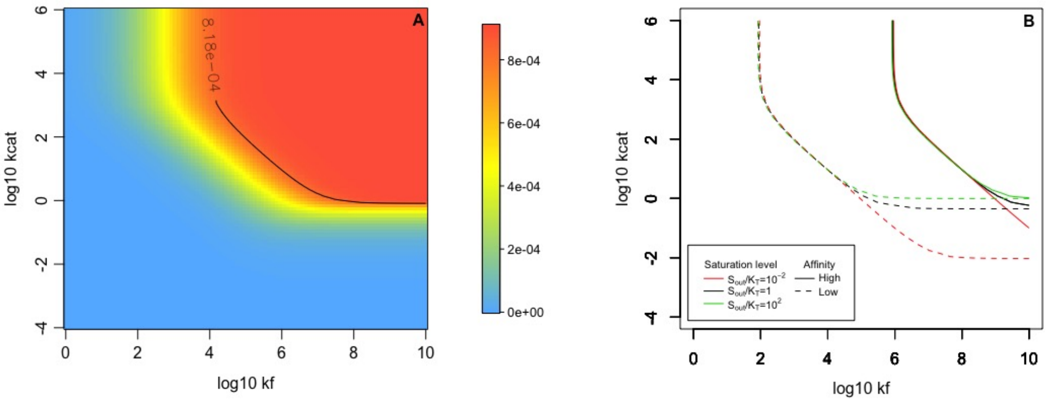


Figure S2: Fitness landscapes for the first enzymes in pathways when uptake results from FD with sugar-like transporters ($V_{Tm} = 10^{-3} M.s^{-1}$). On the right hand side (A), the absolute flux is shown for a moderate affinity and nearly saturated transporters ($[S_{out}] = 10K_T$, $K_T = 1mM$) to compare with absolute flux resulting from PD of sugars (see Figure S1); the isocline corresponds to 90% of the maximum achievable flux. In (B) – where affinity is either high ($K_T = 10\mu M$) or low ($K_T = 0.1M$) – the influence of the substrate concentration in the environment is represented and proven to have little or no impact also in the case of a high flux, especially when the affinity is high.

for lower levels of flux, saturation only affects the selective pressure acting on $k_{cat}$ when transporters affinity is moderate to low.

### Text S2 Transporters and the general shape of the fitness landscape (under FD)

On Figure S3, the fitness landscape of the upstream enzyme is displayed considering different kinetic parameters of transporters (summarized by the 0.9 isocline in fig. 2A of the article) that correspond to those found in living organisms. In well studied organisms, sugars correspond to the (F,I) cases – low to moderate affinity, high flux – and amino acids and nucleosides more or less match with (A) cases – and to a lower degree, (B) and (D). As discussed in the paper, the maximum flux induced by transporters increases the selective pressure on both $k_{cat}$ and $k_f$ of the first enzyme, while their affinity mainly influences selection on $k_f$.

### Text S3 Fitness landscape of the second enzyme (under FD)

#### Text S3.1 Fitness landscapes with a degradation rate

On Figure S4, concentrations of the first product are represented as functions of the second enzyme parameters, perfectly matching flux landscapes displayed in the article down to the details of the shape. Predictably, it shows that the degradation rate exerts an apparent influence once and only once metabolite concentrations start to rise and their product with the degradation rate approach the level of flux achieved within the pathway (slightly below $10^{-6}M$). Indeed, at steady-state, if the next enzyme is very inefficient, almost exactly the flux of newly formed product is degraded, which means that $\eta_d[P_1]^* \approx \Phi_{P_1}$, yielding $[P_1]^* \approx \Phi_{P_1}/\eta_d$.

Equilibrium concentrations for sugars are higher for similar degradation rates because the influx is larger (see Figure S5) - the increase being proportional to that of the flux. If the absolute concentration – instead of the relative cost – matters then the constant $\eta_d$ may be higher for sugars (thereby reaching biologically relevant metabolite concentrations).

Degradation rates modulate the fitness landscapes of the second enzyme. One remarkable phenomenon here is that the landscape does not depend much on the efficiency of the first enzyme : this is especially true for $k_f$, whose selective pressure is mostly insensitive to the first enzyme efficiency (or the level of flux, more generally). For a system where a specific flux of metabolite $\Phi$ can either be converted through a chemical reaction with a dedicated enzyme or degraded, it can be shown that the relative loss of fitness incurred by a change in $k_f$ does not depend on $\Phi$ when $\Phi << k_{cat}[E_{tot}]$

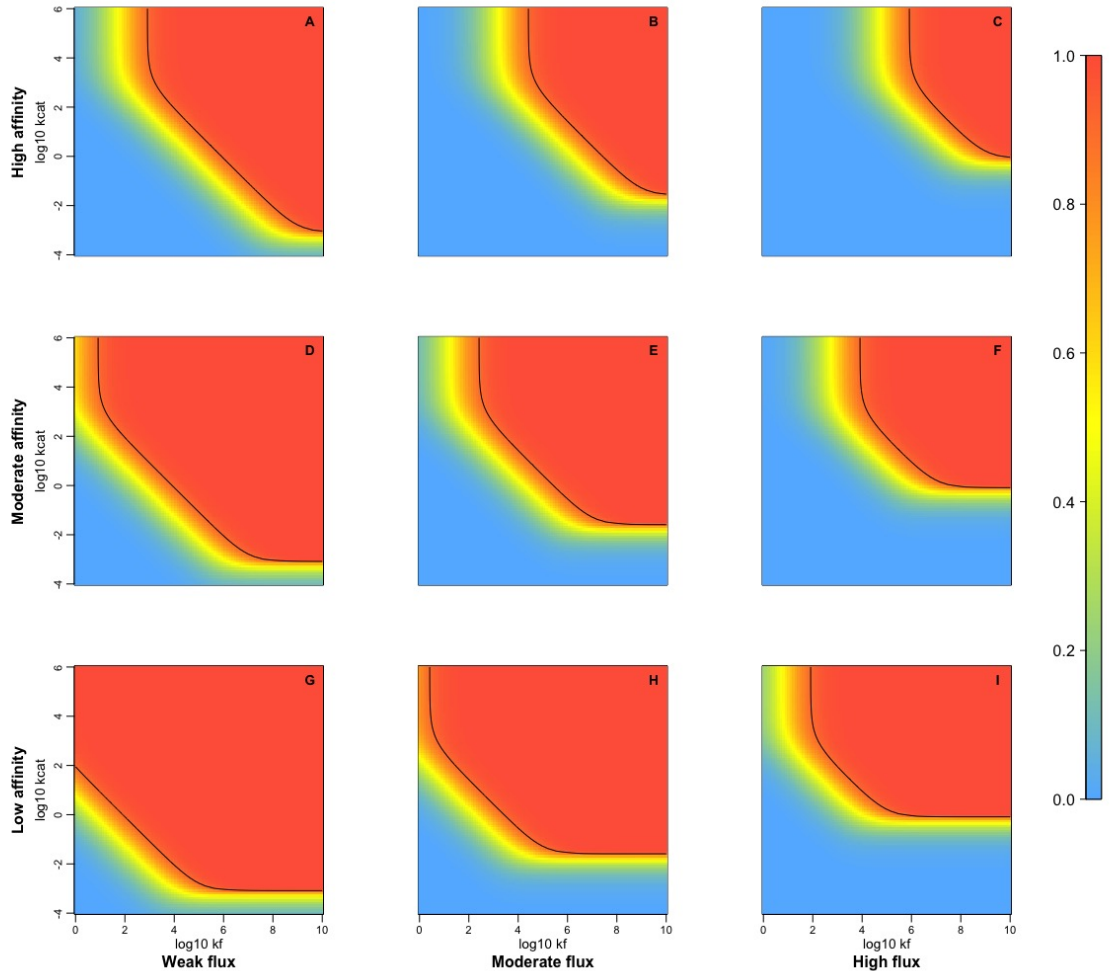


Figure S3: Both the affinity and rate of a transporter have an impact on the (normalized) flux landscape for upstream enzymes, the black isocline (corresponding to 0.9) delineates the fitness plateau. Each plot represents the landscape obtained with a pair of values for transporters affinity $K_T$ and saturation $V_{Tm}$. Moving one step to the right means that $V_{Tm}$ increases by 1.5 orders of magnitude – from $10^{-6}$ (low flux) to $10^{-3} M.s^{-1}$ (high) – and one step up means that $K_T$ decreases by 2 orders of magnitude, starting at $10^{-1} M$ (low affinity). Increasing $K_T$ extends the plateau only towards the left part of the landscape, allowing enzymes with lower $k_f$s on the plateau, whereas decreasing $V_{Tm}$ extends the plateau in both directions. Other parameter values are : $k_r = 1000/s$, $[E_{tot}] = 1mM$ and $[S_{out}] = 10 \times K_T$.

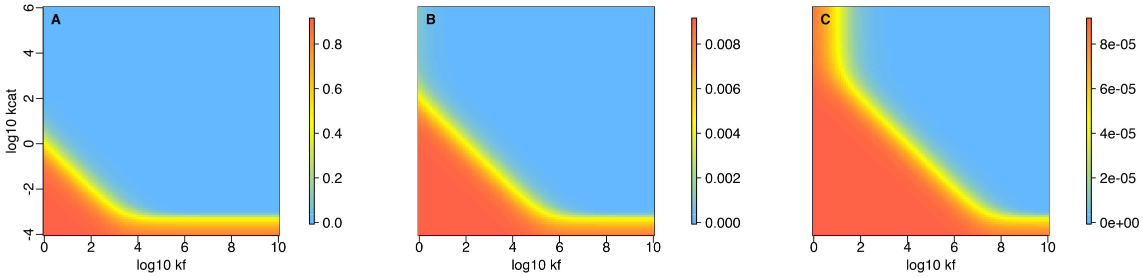


Figure S4: Concentrations (unit: M) of the product of the first reaction for three degradation rates are plotted as functions of second enzyme kinetic parameters, for the model case of nearly saturated amino-acids like transporters ($V_{Tm} = 1\mu M/s$, $K_T = 50\mu M$ and $[S_{\text{out}}] = 10K_T$). The first enzyme is assumed to be perfect and highly concentrated ($k_f = 10^{10} M^{-1}s^{-1}$, $k_{cat} = 10^6 s^{-1}$, $k_r = 10^3 s^{-1}$, $[E_{tot}] = 10^{-3} M$). In (A), the degradation rate is set to $\eta_d = 10^{-6} s^{-1}$ and limits the intracellular concentration of product to $[P_1] = 1M$, while in (B) and (C), these values are respectively $\eta_d = 10^{-4} s^{-1}$ and $10^{-2} s^{-1}$ limiting it to $[P_1] = 10^{-2} M$ and $[P_1] = 10^{-4} M$.

(see Mathematical appendix for the proof, where $V_m = k_{cat}[E_{tot}]$). This explains that the location of the fitness landscape for the second enzyme in this case scarcely depends on the previous enzyme.

The flux of product that the second enzyme needs to process depends on the interplay between

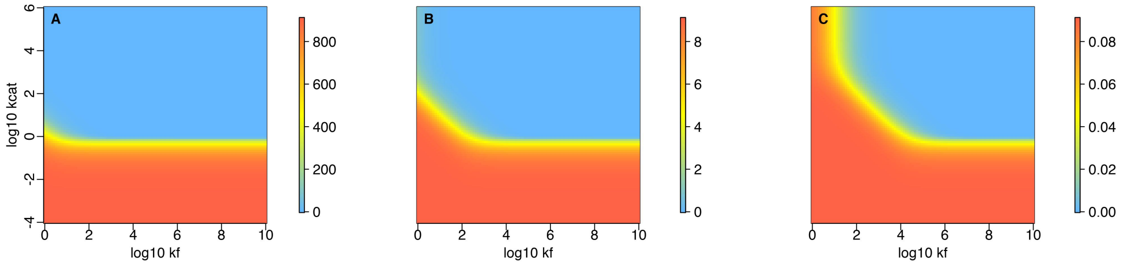


Figure S5: Concentrations (unit: M) of the product of the first reaction for three degradation rates are plotted as functions of second enzyme kinetic parameters, for the model case of nearly saturated sugar like transporters ($V_{Tm} = 1mM/s$, $K_T = 5mM$ and $[S_{\text{out}}] = 10K_T$). The first enzyme is assumed to be perfect and highly concentrated ($k_f = 10^{10}M^{-1}s^{-1}$, $k_{cat} = 10^6 s^{-1}$, $k_r = 10^3 s^{-1}$, $[E_{tot}] = 10^{-3}M$). In (A), the degradation rate is set to $\eta_d = 10^{-6}s^{-1}$ and limits the intracellular concentration of product to $[P_1] = 10^3 M$, while in (B) and (C), these values are respectively $\eta_d = 10^{-4}s^{-1}$ and $10^{-2}s^{-1}$ limiting it to $[P_1] = 10^1 M$ and $[P_1] = 10^{-1}M$. As the first two concentrations (A and B cases) cannot be reached without completely compromising a cell's viability, only the higher one is considered to compare with the first enzyme (fitness landscape in Figure S6) along with two higher degradation rates limiting concentration to $[P_1] = 10^{-2}M$ and $[P_1] = 10^{-4}M$, the same amount than in the case of amino acids with the upper degradation rates.

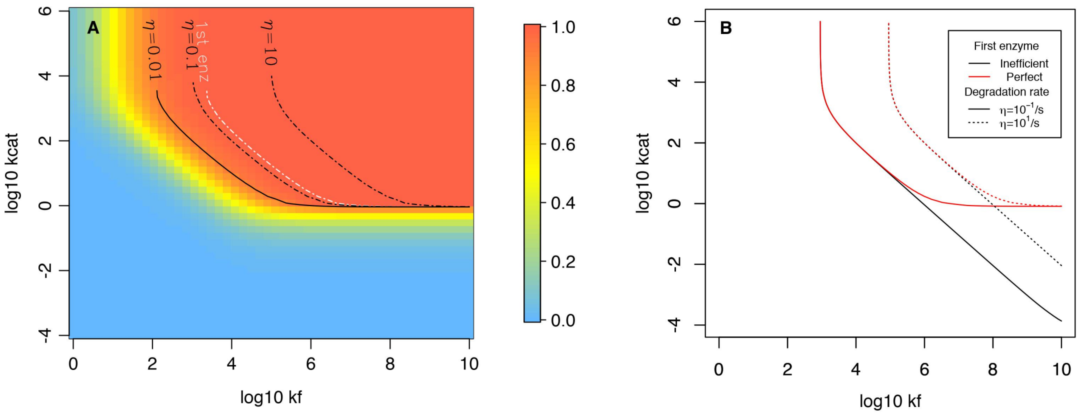


Figure S6: Fitness landscapes for the second enzyme in the context of an irreversible pathway (no backwards flux for the first reaction). These enzymes are involved in a high flux pathway whose pace is driven by $V_{Tm} = 1mM/s$, $K_T = 5mM$ and $[S_{\text{out}}] = 10K_T$. Constant settings for both enzymes also correspond to the model case with $k_r = 1000s^{-1}$ and $[E_{tot}] = 1mM$. (A) shows the fitness landscape for a rather low degradation rate (for the case of sugars) limiting $[P_1]$ to $10^{-1}M$ when the upstream enzyme is perfect and concentrated ($k_f = 10^{10}M^{-1}.s^{-1}$, $k_{cat} = 10^6 s^{-1}$), and constrasts this landscape with that for two higher degradation rates limiting concentration to $[P_1] = 10^{-2}M$ and $[P_1] = 10^{-4}M$. Predictably, the isocline moves toward upper $k_f$s. Besides, contrasting these landscapes with that of the first enzyme shows that they are rather similar. Plateau isoclines drawn in (B) show results for the two higher degradation rates, in the context of an upstream concentrated enzyme being either perfect (see above) or inefficient ($k_f = 10^2 M^{-1}.s^{-1}$, $k_{cat} = 10^{-2}s^{-1}$, $k_r = 10^3 s^{-1}$). Decreasing $\eta_d$ still makes the cell tolerant to higher concentrations of intermediate metabolites (the product of the first reaction), while the efficiency of the first enzyme only influences the pressure on $k_{cat}$ if high intracellular concentrations are not toxic.

the first intracellular enzyme, the degradation rate and transmembrane transporters. To test the selective influence of the latter on the second enzyme, we plotted the landscape of the second enzyme for different degradation rates and first enzyme efficiencies when the pathway is initiated by sugar-like transporters, showing that the processes studied follow similar trends than with transporters involved in lower fluxes (see Figure S6 for sugars, to contrast with landscapes for lower fluxes found in Fig. 3 of the article).

#### Text S3.2 Fitness landscapes with a non-linear toxicity function

So far, we have considered that the flux was decreased due to the loss of product mainly arising from non-specific activities. But fitness may also be impacted because excessive concentrations in one or few metabolites disrupt other pathways or produce damaged metabolites (for example through

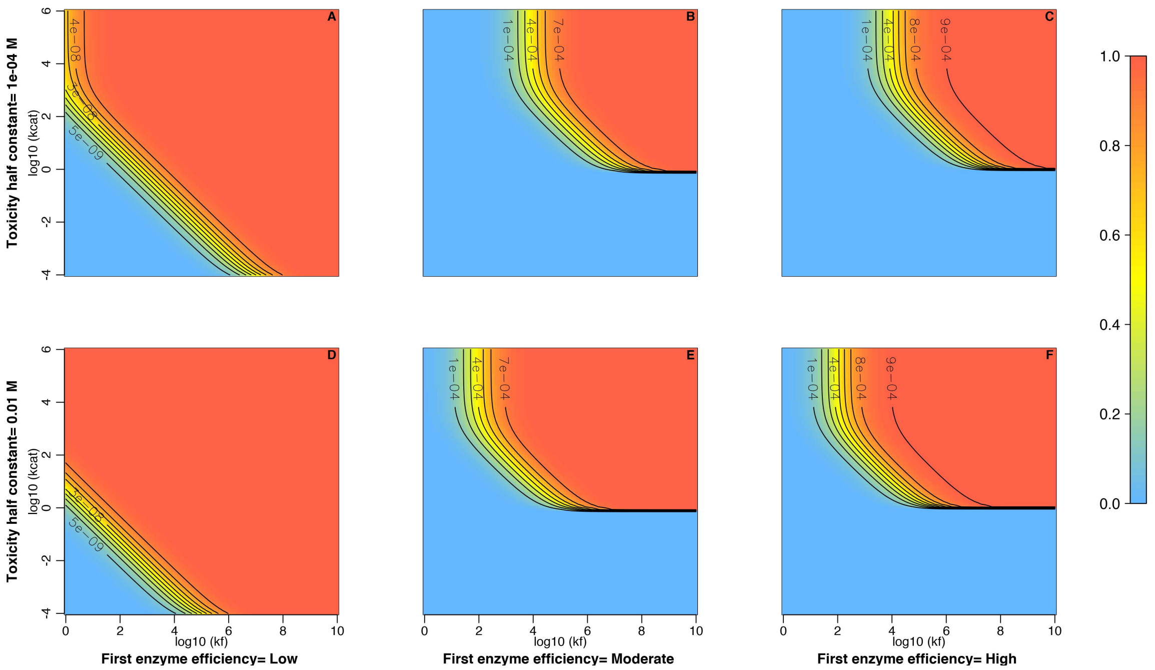


Figure S7: Fitness landscapes for the second enzyme in a pathway when different metabolite toxicity $T$ and first enzyme efficiencies (low : $k_f = 10^2 M^{-1}s^{-1}$, $k_{cat} = 10^{-2}s^{-1}$; moderate : $k_f = 10^5 M^{-1}s^{-1}$, $k_{cat} = 10^1 s^{-1}$; high : $k_f = 10^{10} M^{-1}s^{-1}$, $k_{cat} = 10^6 s^{-1}$) are considered. The case presented here is that of sugar-like transporters ($V_{Tm} = 1mMs^{-1}$ and $K_T = 5mM$). Note that other kinetic parameters are still $[E_{tot}] = 10^{-3}M$, $k_r = 10^3 s^{-1}$ and no reaction reversibility $K_{eq} = 0$. As a bad first enzyme diminishes the flux by several orders of magnitude, the fitness landscape of the second enzyme already flattens for low kinetic values and the influence of toxicity is marginal. On the contrary, the moderately and highly efficient first enzymes give rise to similar fitness landscapes for the second enzyme because they generate relatively similar levels of flux : this is true for both toxicity constants (see Figure S8 for a direct comparison based on relative fitness isoclines), with a linear relationship between the toxicity concentration and the location of the plateau. The fact that the location of the plateau is only determined by the amount of metabolite an enzyme has to process and the cellular tolerance to high concentrations indicates that it applies for any enzyme downstream the first two ones, as expected.

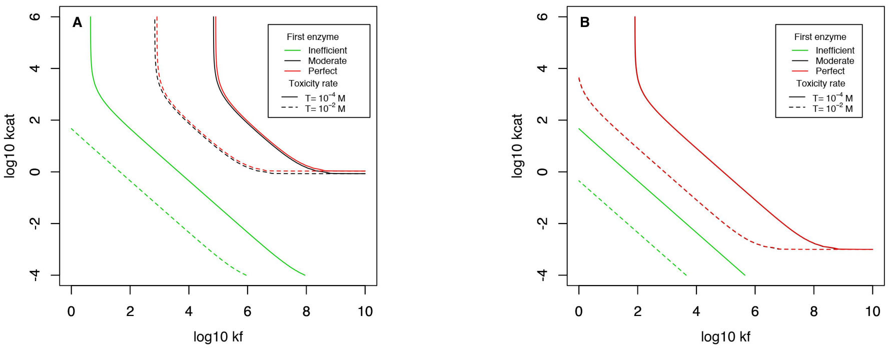


Figure S8: Fitness landscapes (depicted by the 0.9 isoclines) for the second enzyme in a pathway when different metabolite toxicities $T$ and first enzyme efficiencies (low : $k_f = 10^2 M^{-1}s^{-1}$,$k_{cat} = 10^{-2}s^{-1}$; moderate : $k_f = 10^5 M^{-1}s^{-1}$,$k_{cat} = 10^1 s^{-1}$; high : $k_f = 10^{10} M^{-1}s^{-1}$,$k_{cat} = 10^6 s^{-1}$) are considered. Note that other kinetic parameters are still $[E_{tot}] = 10^{-3}M$, $k_r = 10^3 s^{-1}$ and neither reaction reversibility ($K_{eq} \approx 0$) nor degradation rate ($\eta_d = 0$) are considered to show the effect in isolation from the other ones. (A) represents the cases of Figure S7 for which we see that the isoclines showing the level of flux were partly misguiding since the fitness landscapes are in fact complletely superimposable for the cases of a moderately and highly efficient first enzyme. The exact same pattern is found on (B) which represents the case of amino acids like transporters ($V_{Tm} = 10^{-6}M$, $K_T = 50\mu M$). In any case, toxicity proved to display a similar influence, although the shift depends on the level of the flux as shown in Fig. S7. Still, the influence of the first enzyme is different with this form of toxicity than with a linear degradation rate, because the level of flux henceforth matters, which was mostly not the case previously for $k_f$ (see text for the demonstration).

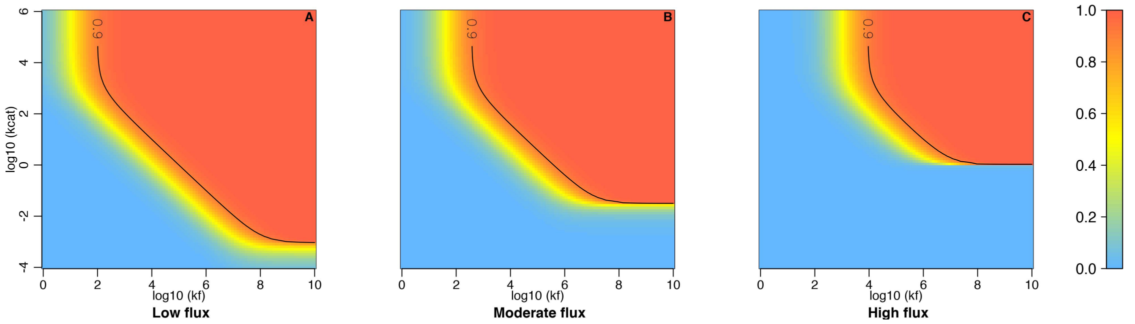


Figure S9: Fitness landscapes of any enzyme for different levels of flux (Low : $\Phi = 10^{-6} M.s^{-1}$; Moderate : $\Phi = 10^{-4.5} M.s^{-1}$; High : $\Phi = 10^{-3} M.s^{-1}$, with a moderate toxicity for the intermediate metabolite ($T = 10^{-3}$) and a moderately high degradation rate $\eta_d = 10^{-2} s^{-1}$. Note that other kinetic parameters are still $[E_{tot}] = 10^{-3} M$, $k_r = 10^3 s^{-1}$ and no reaction reversibility $K_{eq} \approx 0$. One can clearly see that for enzymes involved in non-reversible pathways, the flux is the main driver of the fitness landscape on which an enzyme evolve when metabolite toxicity is accounted for.

promiscuous processes), both phenomena being largely documented (see references in the article). To test the influence that toxicity may have on fitness, we set the absolute fitness to $f = \Phi(1 - \frac{[P_1]}{[P_1]+T})$ (Clark, 1991), such that it results from both the flux of final products (the product formed by the second reaction) and a sigmoid influence dependent on the concentration of the first product $P_1$ (which is processed by the second enzyme to produce the final product). $T$ acts as a threshold concentration which, when getting approached, significantly diminishes fitness. At first approximation, toxicity yields the same effect as the degradation rate (see Figure S8), but in this context, the proximity to the concentration threshold depends on the flux of substrate provided by the reactions upwards in the metabolic pathway. Accordingly, an inefficient first enzyme releases the selection pressure (in green in Figure S8), but not a moderately efficient or perfect first enzyme (see Figure S7).

To gain a better sense of – and generalize – this flux-dependent selection process, we drew fitness landscapes where a fixed supply of substrate is added continously, processed by a perfect enzyme $E_n$ which produces a product $P_n$ that in turn, can eventually be processed by a following enzyme $E_{n+1}$. Toxicity and degradation were set to moderate values, but these parameters do not qualitatively impact the results. It is straightforward on Figure S9 that the flux proportionally increases the selective pressure on enzyme kinetic parameters (increasing the flux by one order of magnitude moves isoclines by one order of magnitude to the upper right). Because we have only considered a given amount of (first or $n^{ieth}$) substrate produced – which can correspond to any reaction at any location within a pathway – the fitness landscape depicted here applies to any enzyme under directional selection to maximize the flux. In fact, the enzyme needs even not be involved in a pathway directly initiated by a transporter and may well be located downstream in branching pathways. It is noteworthy that this dependency between enzymes in a pathway is caused by the non-linear toxicity function; varying the flux under the linear cost function, as described above, would predominantly result in identical fitness landscapes.

### Text S4 Interplay between kinetic parameters

#### Text S4.1 Reversibility and reverse rate influence on fitness landscapes

We have shown in the article that the interplay between kinetic parameters should play a part in the wide variability observed among enzymes, focusing on the effect of inescapable non specific interactions and their possible toxicity (the latter being also detailed in the previous section of the SM). But we have also discussed that physical constraints, the reversibility of reactions in first place, can also explain large differences among enzyme kinetic parameters by competing for the use of a specific substrate – similar to non specific interactions. We here present why reversibility matters and how it impacts the fitness landscape in more details.

Discussion of the direct effect of reversibility can be found in the dedicated section of the paper while the graphical results on which this dicussion is based are presented in Figures S10 for a low

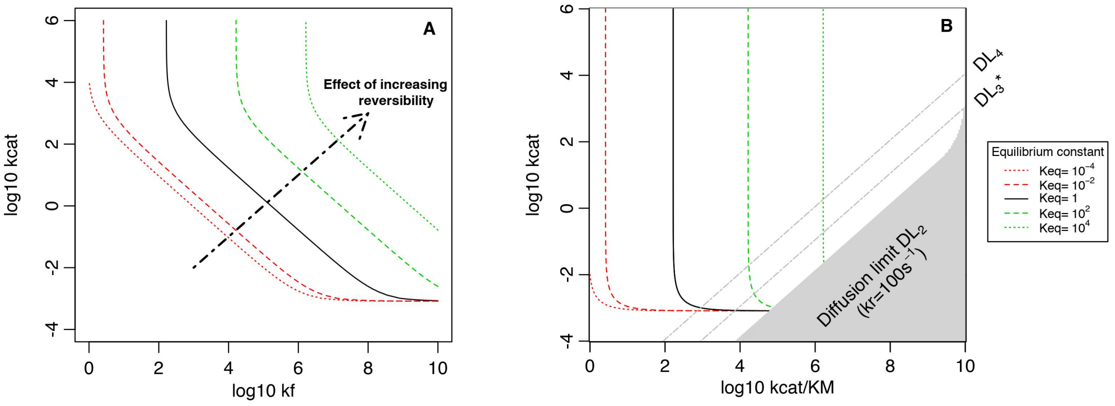


Figure S10: Backwards reaction rates of an enzyme directly upstream have a strong impact on the fitness landscape. Both plots show results of the influence of the reversibility of the first reaction on the fitness landscape of the next enzyme. Parameters are identical as in the model case for amino acids, with $k_r = 10^3 s^{-1}$ and $[E_{tot}] = 1mM$. $K_{eq}$ equals $[S]_{eq}/[P]_{eq} = k_r k_{inh}/k_{\text{cat}} k_f$ (Klipp and Heinrich, 1994) and quantifies the degree of reversibility, a low $K_{eq}$ featuring low reversibility and vice versa. The first enzyme is a nearly perfect forward enzyme, but with $k_f = 10^8 M^{-1}$ and $k_{cat} = 10^4 s^{-1}$ so that we can test for enzymes being even more efficient working in reverse without using abnormal values (eg. association constant overcoming the diffusion limit set to $10^{10} M^{-1} s^{-1}$). Reversibility was equally spread between the two backwards parameters (*e.g.* $K_{eq} = 10^2$ yields $k_r = 10^1 k_{cat}$ and $k_{inh} = 10^1 k_f$), and a low degradation rate was considered ($\eta_d = 10^{-4} s^{-1}$). In (A), results are plotted in the theoretical parameter space ($k_f$, $k_{\text{cat}}$), showing that any increase in reversibility increases the pressure on enzyme kinetics by the same magnitude – except when reactions are highly non-reversible (in red). In (B), the same results are shown in the experimenter parameter space of the second enzyme, showing that there is an increased pressure on $k_{\text{cat}}/K_M$ under higher reversibility. While the plateau is not moved upwards, indicative of a selection on $k_{\text{cat}}$ independent on reversibility when $k_{\text{cat}}/K_M$ is fixed, positive selection for this parameter may still arise due to the diffusion limit that precludes the access to the lowest $k_{\text{cat}}$ at high $k_{\text{cat}}/K_M$. The diffusion limit should play an important role for enzymes with a high dissociation rate $k_r$, as illustrated by the delineation of the diffusion limit (grey area or grey dashed lines) corresponding to several $k_r$ values (eg. $\text{DL}_3$ stands for $k_r = 10^3 s^{-1}$; the star indicates that it is the case represented in (A)).

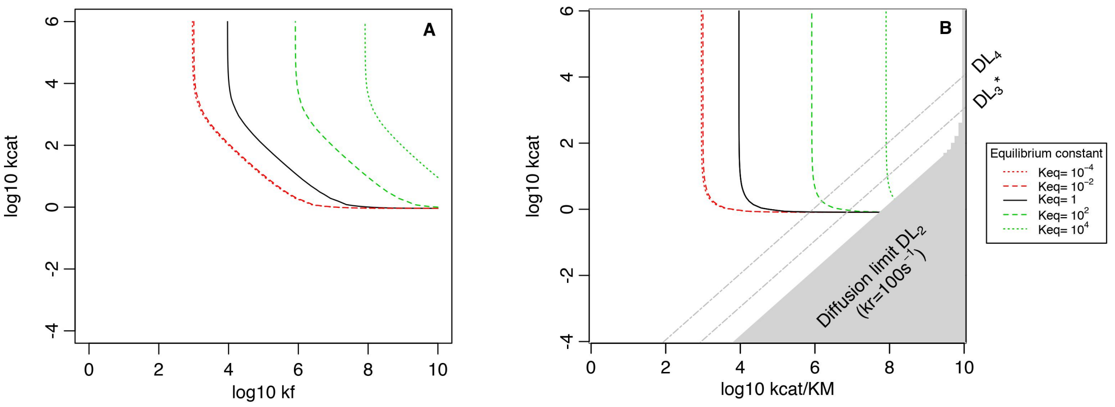


Figure S11: Influence of backwards parameters on fitness landscapes: both plots show results of the influence of the reversibility of the first reaction on the fitness landscape of the following enzyme. Parameters for uptake are $V_{Tm} = 1mM.s^{-1}$ and $K_T = 5mM$, corresponding to the model case for sugars. The settings for enzymes are also the same as the generic ones used previously (highly concentrated enzymes and $k_r = 10^3 s^{-1}$).The first enzyme is a nearly perfect forward enzyme, but with $k_f = 10^8 M^{-1}$ and $k_{cat} = 10^4 s^{-1}$. Reversibility was equally spread between the two backwards parameters (eg. $K_{eq} = 10^2$ yields $k_r = 10^1 k_{cat}$ and $k_{inh} = 10^1 k_f$), and a low degradation rate was considered ($\eta_d = 10^{-4} s^{-1}$). In (A), results are plotted in the theoretical parameter space (made up of $k_f$ and $k_{cat}$), showing that any increase in reversibility increases the pressure on enzyme kinetics by the same magnitude – except when reactions are highly irreversible (in red) – because the first product, instead of accumulating, may thus often react backwards and limit the flow progressing forward. In (B), the same results are shown in the experimenter parameter space of the second enzyme, showing that there is an increased pressure on $k_{cat}/K_M$ (trivially, as the isoclines are displaced towards higher values) but also on $k_{cat}$, first because $k_{cat}$ and $k_{cat}/K_M$ are not independent; second, because the diffusion limit precludes the access to high $k_{cat}/K_M$ for a wider range of $k_{cat}$ values when the pressure on $k_{cat}/K_M$ is higher. This is particularly true when the dissociation rate $k_r$ of the second enzyme is high, as illustrated by the delineation of the diffusion limit (grey area or grey two-dashed lines) corresponding to several $k_r$ values.

flux and S11 for a high flux. Both of them are shown in the theoretical's (A) and experimenter's (B) parameter space to better grasp how basic properties give rise to phenomenological ones. Overall, the equilibrium constant $K_{eq}$ has a rather similar impact on the fitness landscape of the next enzyme than the non-linear toxicity function, with a highly reversible upstream enzyme exerting a selection pressure downstream towards an increase of kinetic parameters.

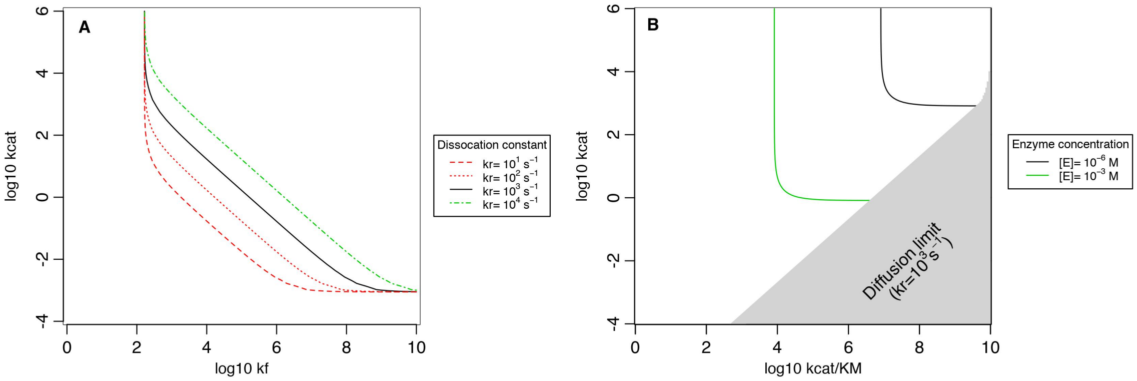


Figure S12: Influence of several kinetic quantities involved in catalytic activity. (A) Influence of the dissociation constant $k_r$ on the joint selective pressure acting on forward parameters $k_{cat}$ and $k_f$ for the model case of the first enzymes involved in the processing of amino acids (see FIG. 1 in main body of article). (B) Influence of enzyme concentration on the pressure acting on forward parameters in the model case for sugars ($V_{Tm} = 1mM.s^{-1}$ and $K_T = 1mM$): see next figure for the effect in the context of amino acids-like pathways, coupled to the influence of reversibility. In this panel, increasing an enzyme concentration is not costly, which is an unrealistic assumption relaxed later.

Increasing reversibility through an increase in $k_r$ (Figure S12-A) increases the selective pressure on both kinetic parameters downstream – except for very low $k_r$ – jointly pushing them towards higher values. When using the "empirical" parameter space (Figure S10-B), we observe that increasing reversibility $K_{eq}$ only selects for higher $k_{\text{cat}}/K_M$ in principle. Yet, the diffusion limit constraint also matters since it removes accessibility to the high $k_{cat}/K_M$, low $k_{cat}$ part of the landscape (see grey area and grey lines on Figure S10-B). As $K_{eq}$ sets the ratio between the four kinetic parameters, the joint evolution of forward and backward rates is determined by the dependency this ratio creates : indeed, higher reversibility comes on average with higher $k_r$s for similar turn-over rates $k_{cat}$, which means that higher reversibility pushes both $k_{cat}$ and $k_{cat}/K_M$ to the upper right corner. On the contrary, when $K_{eq}$ is low, the diffusion limit may vanish from the experimenter's space and opens it up completely. As a consequence, the unbinding rate $k_r$ of an enzyme may eventually fuel a positive codependency between the forward rates $k_{\text{cat}}$ and $k_f$ as the latter is no longer sufficient to ensure a high $k_{\text{cat}}/K_M$ (see Figure S10-B and Figure S11-B of SM for this influence in the theoretical parameter space).

Because optimal concentrations may generally be below $10^{-3}M$ and since the flux is determinant in the drawing of the fitness landscape (especially when accounting for toxicity of metabolites), the influence of epistasis between following enzymes also differs (see Figure S14). This means that when toxicity is the main driver of the selective pressure, the moderate epistasis - between kinetic parameters of neighbouring enzymes - shown with high concentrations (see Figure S8) increases if lower concentrations are assumed.

#### Text S4.2 Enzyme concentration and shape of the fitness landscape

Finally, levels of gene expression can also influence an enzyme's catalytic activity, and as such, high enzyme concentrations can relax the selective pressure acting on enzyme kinetic parameters, whereas very low concentrations require that extreme kinetic efficiencies evolve when the metabolic demand is high like in the case for sugars (see Figure S12B). In this context, the influence of enzyme concentrations is similar for the second enzyme (see Figure S13) such that increasing it comes with a relaxed selective pressure on both $k_{\text{cat}}$ and $k_f$ (and $k_{\text{cat}}/K_M$).

However, we have unreasonably assumed here that gene expression is cost-free, an assumption that we now relax by introducing two well documented costs: 1) the cost of protein production and 2) the cost of (excessive) macromolecular crowding. Therefore, on the one hand $[E_{tot}]$ increases the

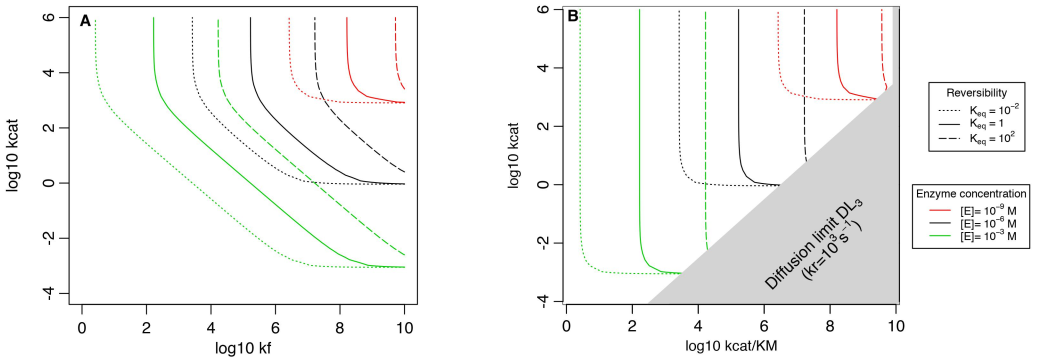


Figure S13: Influence of enzyme concentration on kinetic parameters for the model case of amino acids ($V_{Tm} = 1\mu M.s^{-1}$ and $K_T = 50\mu M$) followed by a nearly perfect enzyme and including a low degradation rate ($\eta_d = 10^{-4}s^{-1}$) for the first product, represented in (A) the theroretical parameter space; and (B) the experimenter's parameter space: the case of downstream enzymes in a pathway. These plots show the joint variability introduced by the reversibility of the previous – the first one here – reaction and the concentration of the enzyme of the focal – the second one here – reaction. General trends are identical than for the first enzyme and reversibility of the previous reaction has the same effect (with the same magnitude) no matter the enzyme concentration. Decreasing enzyme concentration pushes the fitness plateau towards higher values of both $k_{cat}$, $k_f$ and $k_{cat}/K_M$. In this panel, increasing an enzyme concentration is not costly, which is an unrealistic assumption relaxed later.

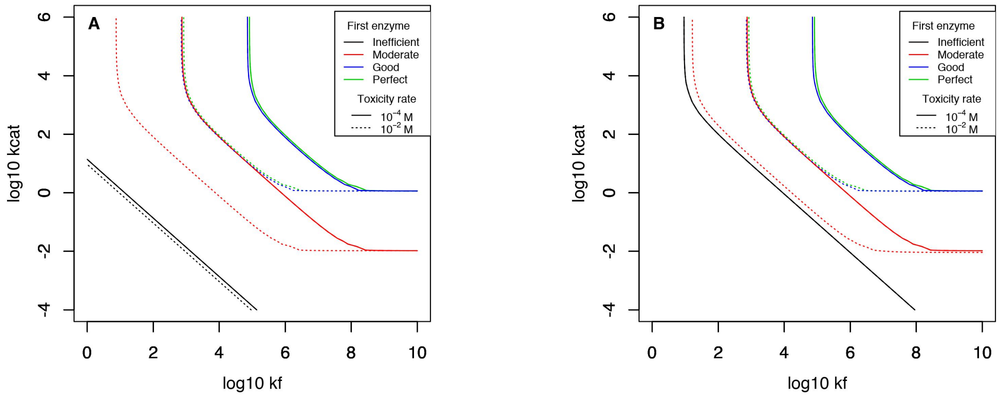


Figure S14: Influence of the first enzyme efficiency on the landscape of a second enzyme involved in the same pathway, considering two different levels of toxicity along a low degradation rate ($\eta_d = 10^{-6}s^{-1}$) in (A) and a moderate one ($\eta_d = 10^{-2}s^{-1}$) in (B). Contrary to preceding figures, the concentration is now set to $[E_{tot}] = 1\mu M$ to see how it may influence epistasis relationships within a pathway. Four first enzyme efficiencies are considered (Inefficient : $k_f = 10^2M^{-1}.s^{-1}$, $k_{cat} = 10^{-2}s^{-1}$; Moderate : $k_f = 10^5M^{-1}.s^{-1}$, $k_{cat} = 10^2s^{-1}$; High : $k_f = 10^7M^{-1}.s^{-1}$, $k_{cat} = 10^4s^{-1}$ ; Perfect : $k_f = 10^{10}M^{-1}.s^{-1}$, $k_{cat} = 10^6s^{-1}$). We see here, that toxicity being a major factor of influence, it changes how the fitness landscape of a second enzyme reacts to different first enzyme efficiencies. This is expected since toxicity dependent lanscapes have been proved to be sensitive to the level of flux. In (B), an inefficient enzyme is more constrained because the higher degradation rate still competes with the second enzyme at relatively low metabolite concentrations.

flux by increasing the enzyme kinetic activity while on the other hand, it hinders diffusion in an exponential manner and reduces the net effect of flux on fitness because part of this flux is diverted to sustain the enzyme concentration (detailed formulas describing these processes can be found in the paper - section Materials and Methods). This interplay gives rise to the fitness landscapes depicted in figure S15 (in which the absolute value of the spread to the maximum fitness is represented on a log-scale such that highest values still correspond to the highest fitnesses), where a crowding limit emerges mechanistically (in grey) for concentrations corresponding to the glass transition (Dill *et al.*, 2011). Besides, the optimum concentration is shifted owing to the cost of production, a phenomenon which is both dependent on the kinetic parameters of the enzymes and the estimate of the cost used.

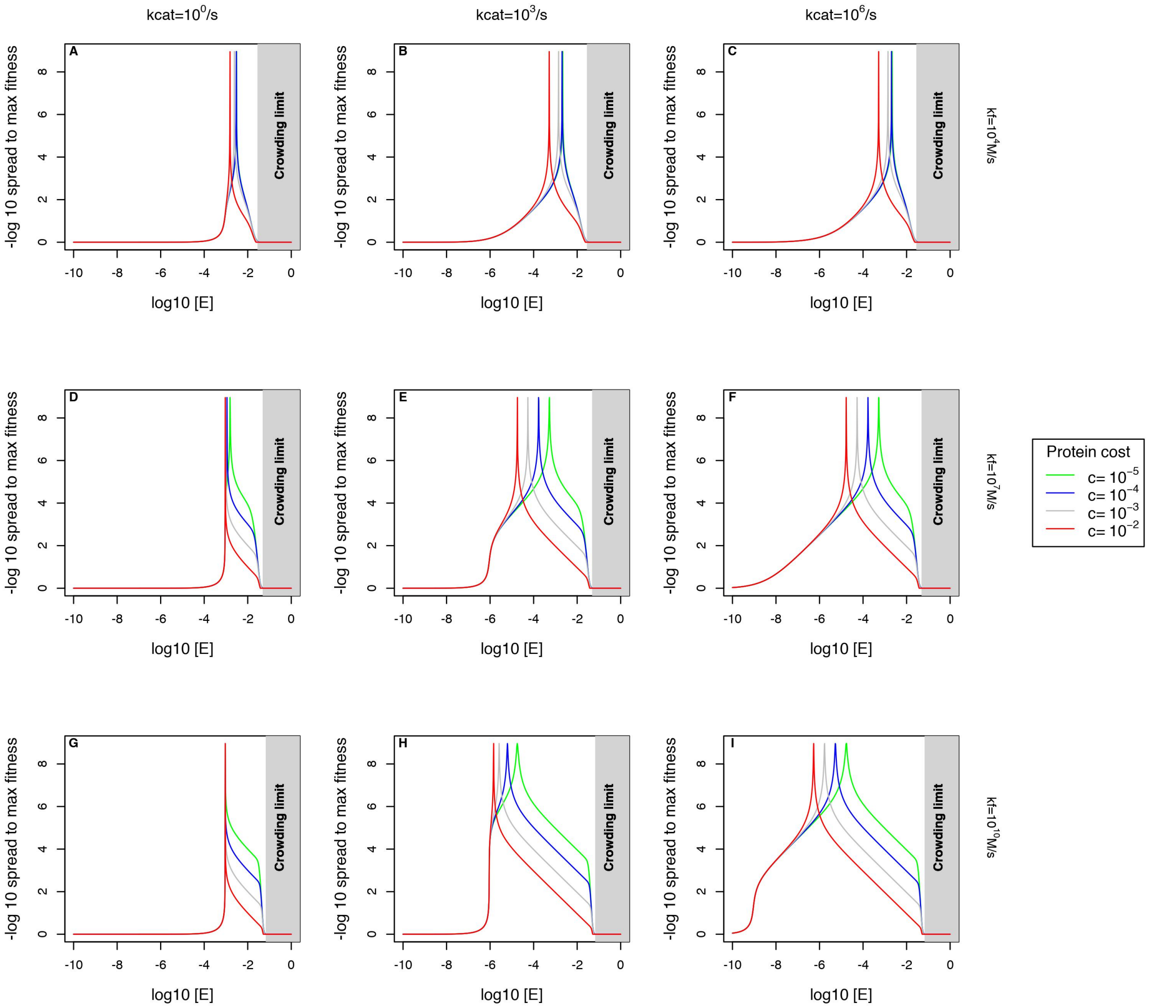


Figure S15: Influence of enzyme concentration on the fitness landscapes for different combinations of enzyme kinetic parameters and different costs of production. The panels represent the absolute value of the $log_{10}$ spread between the maximum fitness and the fitness for the concentration represented on the x-axis, such that a peak corresponds to high fitness. $k_{cat}$ improves from left to right (specific values indicated at the top of each column) and $k_f$ improves from top to bottom (specific values indicated at the right of each row). The grey area delimits the crowding limit where fitness equals zero because the cytoplasm undergoes the glass transition; obviously, because diffusion slows down before reaching this level, fitness starts to plummet for concentrations approximately an order of magnitude below (see the case of a moderately low $k_f$ on the first row, where the cost of expression is not the dominant constraint). In parallel, the protein cost exerts a linear influence with expression, which is especially important when kinetic parameters are higher, as outlined by the cost-dependent spread of the influence of concentration in the lower panels. This means that Natural Selection should oppose the protein burden differently for protein of different costs and that drift should in turn act differently upon enzyme kinetic parameters depending on the values of $N_e$s since the burden differs with these parameters.

### Text S5 Evolutionary simulations

Due to the interplay between drift, mutation and selection, Evolution eventually establishes steady-states in which enzyme kinetic parameters walk randomly – in response to nearly neutral mutations – in a given part of the landscape. To study the mutation-selection-drift balance for enzymes, we first verified that the equilibrium was achieved for each set of parameters detailed in the main body of the article and for each instance of the model (see the top 4 rows of Figure S22 for moderate metabolic demands corresponding to amino-acids/nucleotides for instance, Figure S23 for high metabolic demand corresponding to sugars; Figures S24, S25, S26 for the cases of a low flux when considering mutational correlations; Figures S27, S28, S29 for the cases considering the joint evolution of kinetic constants with enzyme concentrations under different set of assumptions). Watching carefully dynamics, one can notice that clonal interference seems to curb a little the evolutionary process inpopulations with higher $N_e$s, as expected.

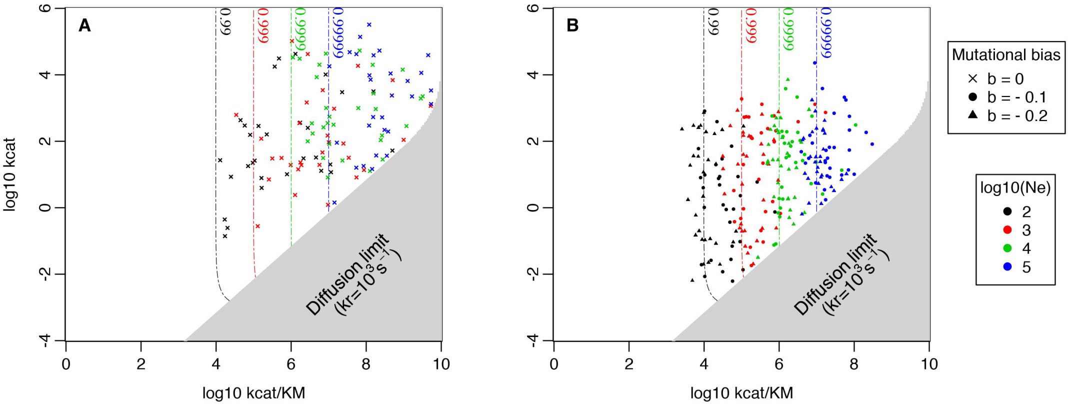


Figure S16: Comparison between evolutionary steady-states and fitness landscapes depicted by isoclines – for the case of a low flux with high affinity ($V_{Tm} = 1\mu Ms^{-1}$ and $K_T = 10^{-5}M$) – under various scenarios: four effective population sizes from $10^2$ to $10^5$ (different colors) and three cases of mutational biases $b$ ($b = 0$ corresponds to the absence of mutational bias). For each scenario, 30 independent simulations were ran, whose outcomes are represented by a dot (per simulation), here in the experimenter parameter space ($k_{\text{cat}}$ and $k_{\text{cat}}/K_M$). Only $k_{\text{cat}}$ and $k_f$ were susceptible to evolve, while $k_r$ was set to $10^3 s^{-1}$ such that the grey part of the parameter space is inaccessible to enzymes due to the diffusion limit. At evolutionary steady-state, enzyme efficiencies evolve on the plateau in any case, each plateau starting according to the strength of drift enzymes cope with. In (A), no mutational bias results in evolutionary outcomes that spread all over the plateau, some reaching very high $k_{\text{cat}}$ and/or $k_{\text{cat}}/K_M$ values, while in (B) enzyme efficiencies stick to their predicted isoclines – under the Nearly Neutral Theory of Evolution (Ohta, 1992) – owing to the over-representation of mutations that diminish efficiency. The stickiness to the isocline is self-evidently positively correlated to the average mutational bias.

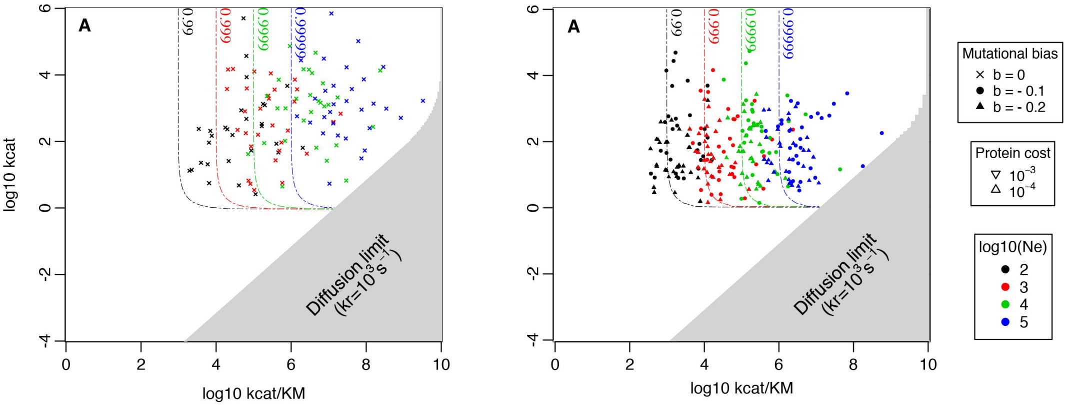


Figure S17: Comparison between evolutionary steady-states and fitness landscapes depicted by isoclines – for the case of a high flux with low affinity ($V_{Tm} = 1mMs^{-1}$ and $K_T = 0.1M$) – under various scenarios: four effective population sizes from $10^2$ to $10^5$ (different colors) and three cases of mutational biases $b$ ($b = 0$ corresponds to the absence of mutational bias). For each scenario, 30 independent simulations were ran, whose outcomes are represented by a dot (per simulation), here in the experimenter parameter space ($k_{\text{cat}}$ and $k_{\text{cat}}/K_M$). Only $k_{\text{cat}}$ and $k_f$ were susceptible to evolve, while $k_r$ was set to $10^3 s^{-1}$ such that the grey part of the parameter space is inaccessible to enzymes due to the diffusion limit. At evolutionary steady-state, enzyme efficiencies evolve on the plateau in any case, each plateau starting according to the strength of drift enzymes cope with. In (A), no mutational bias results in evolutionary outcomes that spread all over the plateau, some reaching very high $k_{\text{cat}}$ and/or $k_{\text{cat}}/K_M$ values, while in (B) enzyme efficiencies stick to their predicted isoclines – under the Nearly Neutral Theory of Evolution (Ohta, 1992) – owing to the over-representation of mutations that diminish efficiency. The stickiness to the isocline is self-evidently positively correlated to the average mutational bias.

#### Text S5.1 Mutation-Selection-Drift balance of kinetic parameters

As explained in the article, fitness at steady-state matches the expectations of the Nearly Neutral Theory (Kimura, 1968; Ohta, 1973, 1992), with average fitnesses aligned with drift barriers (Sung *et al.*, 2012) when mutations are biased towards making enzymes less efficient, and a large nearly neutral area (more fitness variability) when there is no bias at all (see bottom lines of plots in Figures S22 and S23 when no mutational correlations are considered; that in Figures S24, S25, S26 otherwise - in the case of a low flux). No significant difference on the evolutionary trends exists between simulations based on enzymes involved in pathways initiated by transporters with distinct rates $V_{Tm}$. Therefore, when no mutational bias is considered, enzyme kinetic parameters cover the whole nearly neutral plateau that begins when fitness has reached the drift barrier isocline, which stands at $w = 1 - 1/N_e$ relatively to the maximum achievable level (because mutations occuring at this level of fitness cannot provide more than $s = 1/N_e$ of extra fitness). It ensues that enzymes display a wide variability in robustness – as seen in Figures S16 and S17.

Introducing mutational correlations through bivariate gaussian distributions (see Materials and Methods in the paper) did not change qualitatively outcomes of the mutation-selection-drift balance : when an unlikely high trade-off is considered, enzymes hit the drift barrier a little earlier because of the extreme mutational pressure. On the contrary, a yet more unlikely positive mutational correlations pull enzyme parameters up by approximately an order of magnitude on average, merely because positive mutations provide nearly twice the effect they would in the other cases (for instance increasing $k_{cat}$ and $k_f$ by half an order of magnitude is rather similar to increasing either one by an order of magnitude, at least for some combinations of parameters). Expectingly, a positive correlation therefore favours combination of more homogeneous kinetic parameters (compare (C) with (A) and (B) in Figure S18), simply because positive mutations on $k_{cat}/K_M$ are more often correlated to positive mutations on $k_{cat}$.

#### Text S5.2 Mutation-Selection-Drift balance of kinetic parameters with enzyme concentrations

In this section, we report results in which enzyme concentrations are made evolvable quantities subject to the two aforementionned costs (1) crowding; (2) protein production.

Accounting only for (1) results in an increase in variance mainly within effective population size replicates (see Figure S19) and a shift from the expected isocline based on a high concentration : as enzymes can partially compensate for differences in kinetic activities due to kinetic constants of enzymes, it enables enzymes to explore a little wider area for a given effective population size.

Accounting only for the cost of production ((2); see Figure S20) largely increases the variance for a given effective population size because higher concentrations can compensate for lower kinetic parameters as long as they do not overcome a cost threshold set by $N_e$, and higher concentrations can more readily evolve (unbiased mutational effects and no physical ceiling in the model) than higher kinetic parameters, which are again considered to be biased. Depending on the relative cost of expression, an enzyme can evolve towards very low kinetic parameters thanks to extremely high (and unrealistic) concentrations or, on the contrary, evolve towards very high kinetic parameters due to the early onset of the protein burden. Because the level of expression under the mutation-selection-drift balance is determined by $N_e$, results are still largely dependent on this factor (in fact, even more than when enzyme concentration is set - see paper for discussion). None of these two independent constraints considered in isolation realistically catch the overall protein burden. In Figure S21, results are shown including both of these costs (like in the article) but for the 4 different costs. Depending on $N_e$ and the cost of production, the two constraints are now affecting the evolutionary outcomes: when the cost is unlikely low, outcomes resemble that of crowding alone while when it is high, they resemble outcomes found when considering protein production alone. Between theses two cases, a mixture of the two constraints affects enzyme steay-states and is discussed in the article since they correspond to the more likely ones.

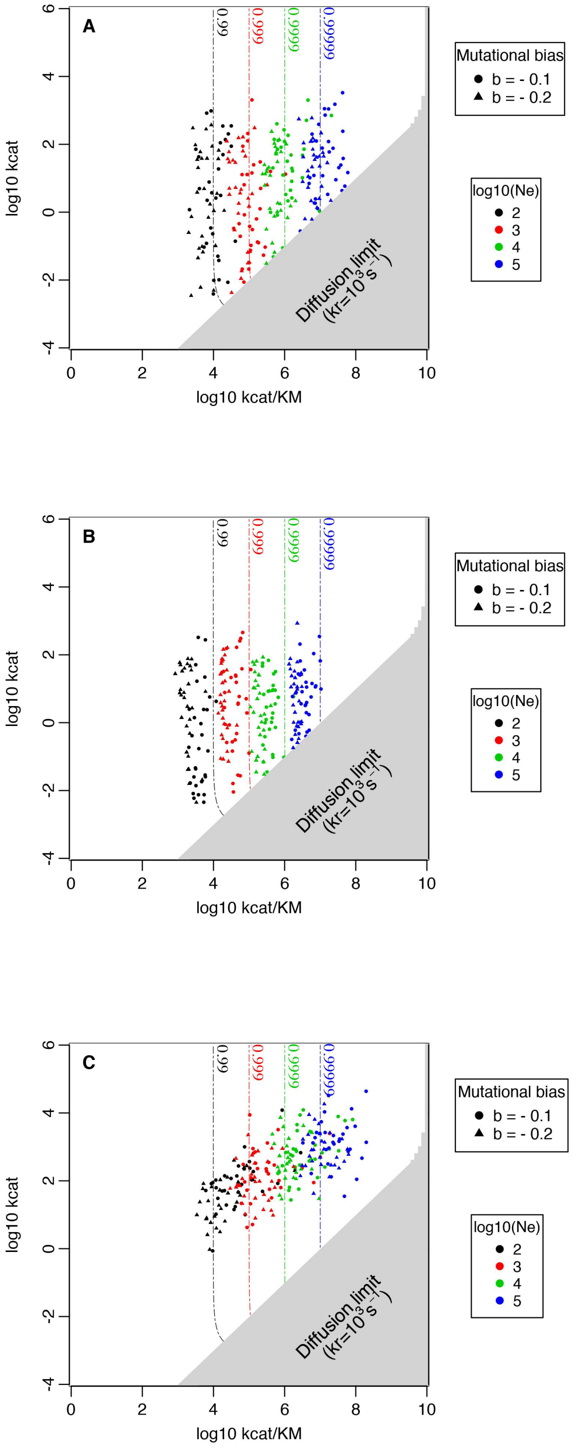


Figure S18: Comparison between evolutionary steady-states and fitness landscapes depicted by isoclines – for the case of correlated mutations and a low flux with high affinity ($V_{Tm} = 1 \mu M s^{-1}$ and $K_T = 10 \mu M$) – under various scenarios: four effective population sizes from $10^2$ to $10^5$ (different colors) and two cases of mutational biases $b$. For each scenario, 30 independent simulations were ran, whose outcomes are represented by a dot (per simulation), here in the experimenter parameter space ($k_{\text{cat}}$ and $k_{\text{cat}}/K_M$). Only $k_{\text{cat}}$ and $k_f$ were susceptible to evolve, while $k_r$ was set to $10^3 s^{-1}$ such that the grey part of the parameter space is inaccessible to enzymes due to the diffusion limit. In (A), a moderate negative correlation ($\rho = -0.5$) is considered, that does not change much the outcomes from the case of no correlation at all, while a higher negative correlation ($\rho = -0.8$) and a positive correlation ($\rho = 0.5$) did slightly change enzyme kinetics at steady-state. Indeed, enzyme efficiencies at the mutation-selection-drift balance stands in the vicinity of their respective drift barrier, albeit negative correlations can pull enzyme fitnesses more or less an order of magnitude down depending on their strength (and the other way around for a positive correlation). In (C), enzymes are also pulled away from the diffusion limit because increases of $k_{cat}/K_M$ are much more often corerlated to increased $k_{cat}$ due to the correlation. .

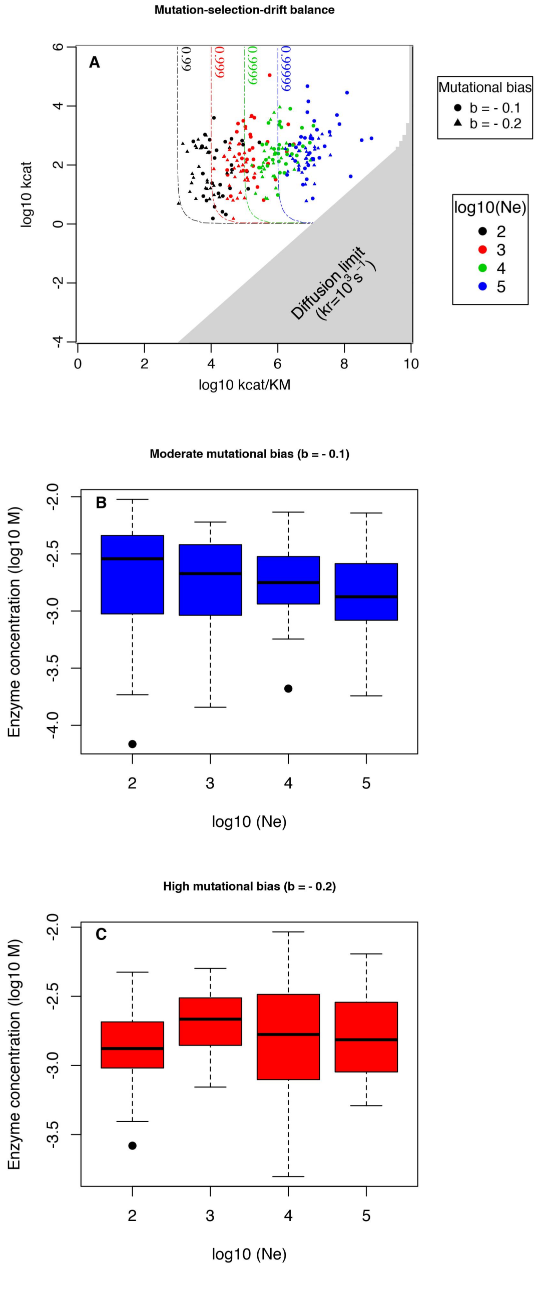


Figure S19: Evolutionary outcomes for enzyme fitnesses and kinetic constants in the case of a high flux with low affinity ($V_{Tm} = 10^{-3}M$; $K_T = 1mM$) where the cost of expression includes only macromolecular crowding effects. Again, 30 simulations were ran for each set of parameter and 2 mutational biases were considered for enzyme kinetic constants, while levels of expression were not subject to mutational bias on the course of their evolution. The first plot (A) shows how enzyme efficiencies spread in the landscape when the mutation-selection-drift balance has established. Values are slightly higher than in the case of fixed enzyme concentrations although concentrations lie on average at sligthly upper levels (see (B) and (C), with no influence of the mutational bias). This means that fighting against this cost comes with a slightly higher directional selective pressure.

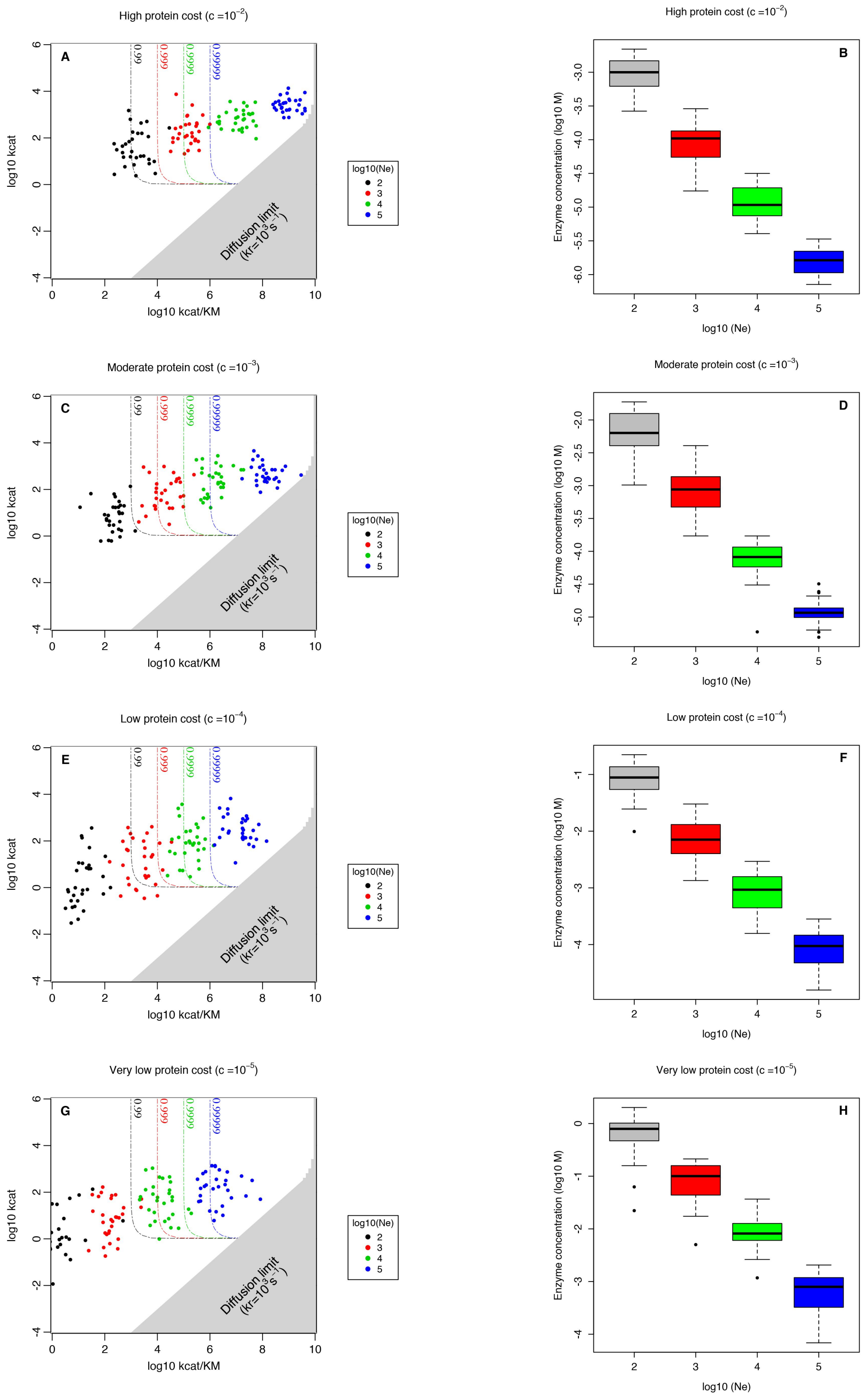


Figure S20: Evolutionary outcomes for enzyme fitnesses and kinetic constants in the case of a high flux with low affinity ($V_{Tm} = 10^{-3}M$; $K_T = 1mM$) when the cost of expression includes only the cost of production. Again, 30 simulations were ran for each set of parameter and a mutational bias (b=-0.2) were considered for enzyme kinetic constants, while 4 costs of production (scaled with regard to a high concentration of $10^{-3}M$) were considered, each one corresponding to a row above. Plots on the left represent the spread of enzyme kinetic constants while those on the right show the fitness reached – both under the mutation-selection-drift balance. As a general rule, enzyme concentrations allow for a little wider exploration in the vicinity of the expected $N_e$-dependent isocline (here that for $[E_{tot}] = 1mM$). Besides, the cost increases the difference between $N_e$s because Natural Selection now more or less actively select against the protein burden in relation to the strength of drift. Note also that absurd concentrations can be achieved since there is no deleterious effects of crowding (see below for the combination of the two costs).

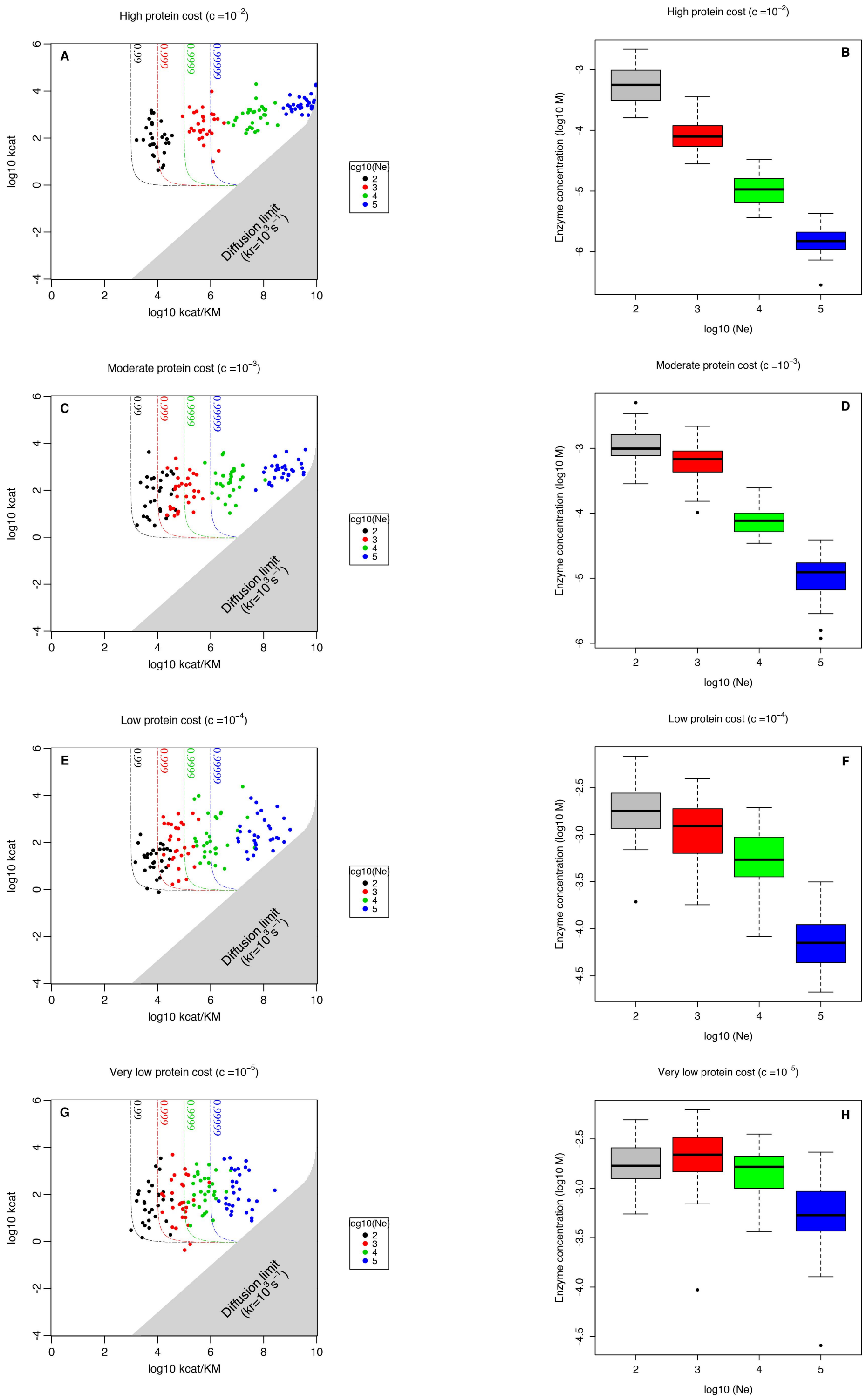


Figure S21: Evolutionary outcomes for enzyme fitnesses and kinetic constants in the case of a high flux with low affinity ($V_{Tm} = 10^{-3}M$; $K_T = 1mM$) when the cost of expression includes both macromolecular crowding and the expenses of production. Again, 30 simulations were ran for each set of parameter and a mutational bias (b=-0.2) were considered for enzyme kinetic constants, while 4 costs of production (scaled with regard to a high concentration of $10^{-3}M$) were considered, each one corresponding to a row above. Plots on the left represent the spread of enzyme kinetic constants while those on the right show the fitness reached – both under the mutation-selection-drift balance. Now that crowding also limits concentration, the outcomes become more or less sensitive to the cost of enzyme production: for an unlikely very low cost, resultts are much alike those obtained with crowding alone. However, when assuming higher costs, outcomes depend on a balance between these two influences, which tend to increase the selective screening for higher $N_e$. As a consequence, enzymes are pushed towards higher efficiencies than when considering high enzyme concentrations, which is all the more true with higher $N_e$s.

### Text S6 Influence of noise in gene expression on the selective pressure acting on enzyme concentrations

One explanation that can account for the discrepancy between these clear-cut results and empirical data is noise in gene expression, whose influence is also manifold as shown by (Wang and Zhang, 2011). In the competition for resources, a cell needs to possess exactly the right amount of enzymes for a given set of kinetic parameters. To ensure that this happens, a cell has to possess continously at least one transcript of a specific enzyme. Since typical prokaryotic cells range around $V_{cell} = 1\mu m^3$, for a prokaryotic cell to have at least one transcript yields a concentration $[RNAm] = \frac{1/(6.02\times10^{23})}{10^{-15}} \approx 1nM$. Making the conservative hypothesis of a 1:100 ratio between transcripts and proteins, this yield a noisy concentration threshold around $[E_{tot}] \approx 0.1\mu M$ for this kind of cells. This is a highly conservative estimate of this threshold, for 1 transcript on average is far from sufficient to alleviate noisiness and even less to tune the kinetic activity with the needed subtelty. As a consequence, the threshold should be one or two order of magnitude higher and typically stands around $1 - 10\mu M$ for average prokaryotic cells, therefore precluding the access to low concentrations in prokaryotes in a volume dependent manner, with smaller prokaryotes being even more constrained than the above estimate. Small cells are thus subject to a complex balancing selection acting on their protein content that needs to be adressed carefully to better understand how enzyme concentrations are tuned, but in Prokaryotes at the least, concentrations are restricted within a narrow range which cannot explain most of the observed variability.

### Text S7 Mathematical and computational appendix

#### General description of nutrient driven pathways - analytical solution for the first enzyme

The model describing the metabolism (through the first enzyme) inside the cell relies on the following system of equations:

$$\begin{cases} \dfrac{d[ES]}{dt} = k_f.[E].[S] - (k_{cat} + k_r).[ES], & \text{(S1a)} \\ \dfrac{d[P]}{dt} = k_{cat}.[ES], & \text{(S1b)} \\ [E_{tot}] = [E] + [ES], & \text{(S1c)} \end{cases}$$

where $[E]$, $[S]$ and $[ES]$ denote the respective cytoplasmic concentrations of the enzyme, substrate, and of their complexes. We assume that the system can reach an equilibrium obeying $\frac{d[ES]}{dt}\Big|_{[ES]=[ES]^*} = 0$, resulting in the following equation:

$$[ES]^* = \frac{k_f[S]^*}{k_r + k_{cat} + k_f[S]^*}.[E_{tot}], \tag{S2}$$

where the equilibrium concentration of the substrate within the cell $[S]$ depends on the uptake mechanism considered, as described in the next two sections.

##### Uptake by passive diffusion

Passive diffusion occurs when molecules cross the membrane without specifically binding any protein. This well-known process can be modelled through Fick's diffusion law (Fick, 1855; Overton, 1899; Meyer, 1899) that takes into account the surface area of a cell $S_c$, its permeability $P$ and the outside-inside concentration gradient. Ignoring cell growth – or, similarly, assuming growth to be far slower than diffusion – this equation can be written as:

$$\frac{d[S]}{dt} = \frac{P.S_c}{V_c}.([S_{\text{out}}] - [S]), \tag{S3}$$

recalling that $[S]$ is the intracellular substrate concentration, and with $V_c$ the volume of the cytoplasm. Assuming a large surrounding environment where the external concentration of the substrate $[S_{\text{out}}]$ remains constant[1] and an homogeneous distribution of the substrate inside the cytoplasm, the dynamics of $[S]$ should obey the following differential equation:

$$\frac{d[S]}{dt} = \frac{P.S_c}{V_c}.([S_{\text{out}}] - [S]) - k_f.[E].[S] + k_r.[ES], \tag{S4}$$

such that the equilibrium value $[S]^*$ (where $\frac{d[S]}{dt} = 0$) obeys the equation:

$$\frac{P.S_c}{V_c}.([S_{\text{out}}] - [S]^*) + k_r\frac{k_f[S]^*[E_{tot}]}{k_r + k_{cat} + k_f[S]^*} - k_f[S]^*\frac{(k_r + k_{cat})[E_{tot}]}{k_r + k_{cat} + k_f[S]^*} = 0.$$

which can be written:

$$-Pk_f\frac{S_c}{V_c}([S]^*)^2 + (P\frac{S_c}{V_c}([S_{out}]k_f - (k_r + k_{cat})) - k_{cat}k_f[E_{tot}])[S]^* + P\frac{S_c}{V_c}[S_{out}](k_r + k_{cat}) = 0 \tag{S5}$$

Since a concentration cannot be negative and this quadratic equation has a single positive root, we obtain a single equilibrium value $[S]^*$ under passive diffusion, whose calculation is straightforward.

[1] A depleting environment would bring back to a quasi steady state situation in which the substrate concentration decreases although on a much slower timescale than that at which happen chemical reactions and transport.

##### Uptake by facilitated diffusion

Assuming that transport is bidirectional and symmetric, FD obeys Michaelis Menten-like kinetics (Kotyk, 1967) that can be modelled through Briggs-Haldane equations (Briggs and Haldane, 1925):

$$\frac{d[S]}{dt} = V_{Tm}.\frac{[S_{\text{out}}] - [S]}{K_T + ([S_{\text{out}}] + [S]) + \alpha.\frac{[S_{\text{out}}][S]}{K_T}} \tag{S6}$$

with:

$$\begin{cases} V_{Tm}\text{: the maximum rate of a given carrier protein;} \\ K_T\text{: a constant inversely proportional to the transporter efficiency;} \\ \alpha\text{ : the Kotyk interactive constant capturing the disequilibrium between bound and free transporters.} \end{cases}$$

Along the concentration gradient, the Kotyk interactive constant $\alpha$ Kotyk (1967) also brings by something new because transporters saturation (*i.e. outwards flow*) can no longer be overlooked (Teusink *et al.*, 1998; Bosdriesz *et al.*, 2018). By construction, $\alpha$ cannot exceed 1 and is apparently close to this upper limit for sugars (*e.g.* $\alpha = 0.91$ for glucose Kotyk (1967)) so we set $\alpha = 1$ by default in this study. With similar assumptions to the PD derivation above regarding the external environment, the inward net flow of the substrate is given by:

$$\frac{d[S]}{dt} = V_{Tm}.\frac{[S_{\text{out}}] - [S]}{K_T + ([S_{\text{out}}] + [S]) + \alpha\frac{[S_{\text{out}}][S]}{K_T}} + k_r[ES] - k_f[S][E],$$

which at steady-state yields a unique positive solution $[S]^*$ because the system can be summarized by equations (8) and (9) found in the Materials and Methods of the main body of the article, after following the same steps than for PD.

#### Flow in metabolic pathways

The flow of product is obtained by substituting $[S]$ in equation (S2) with its steady-state value under either passive or facilitated diffusion, and merging with equation (S1b):

$$\frac{d[P]}{dt} = \frac{k_{cat}k_f[S]^*}{k_r + k_{cat} + k_f[S]^*}.[E_{tot}]. \tag{S7}$$

#### Influence of the metabolite influx and the degradation rate for the fitness landscape of an enzyme

Let $\Phi$ be the influx of a specific metabolite $M$ (which can be the product of the previous enzyme). Such a process - where an enzme competes against degradation to produce its product - can be described by the following expression of flux conservation:

$$\Phi = V_m\frac{[M]}{K_M + [M]} + \eta_d[M] \tag{S8}$$

Note that this equation is similar to that of (S7) where $\frac{d[P]}{dt} = \Phi$, $k_{cat}[E_{tot}] = V_m$ and $\frac{k_r+k_{cat}}{k_f} = K_M$, with the difference that a degradation rate has been added. Rearranging the terms is straightforward and yields a quadratic equation whose only valid solution is:

$$[M]^* = \frac{b(-1 + (1 - \frac{4ac}{b^2})^{1/2})}{2a}, \tag{S9}$$

where:

$$\begin{cases} a = \eta_d \\ b = V_m + \eta_d K_M - \Phi \\ c = -\Phi K_M \end{cases}$$

When $\Phi << V_m$ one can write:

$$\frac{ac}{b^2} \approx \frac{\eta_d \Phi K_M}{Vm^2 + 2\eta_d K_M V_m + (\eta_d K_M)^2}$$

which yields:

$$\frac{ac}{b^2} \approx \begin{cases} \frac{\eta_d \Phi K_M}{Vm^2}, \text{if } \eta_d K_M << V_m \\ \frac{\Phi}{4Vm}, \text{if } \eta_d K_M \approx V_m \\ \frac{\Phi}{\eta_d K_M}, \text{otherwise.} \end{cases}$$

Therefore, if $\Phi << V_m$, one has $\frac{ac}{b^2} << 1$ in any case, such that it is always possible to approximate $[M]^*$ through its first order Taylor series expansion, which yields $[M]^* = \frac{b(-1+(1-2\frac{ac}{b^2}))}{2a} = \frac{-c}{b}$. This means that $[M]^* = \frac{\Phi K_M}{V_m + \eta_d K_M - \Phi}$, and, as we are interested in the case where $\Phi << V_m$, $[M]^* \approx \frac{\Phi K_M}{V_m + \eta_d K_M}$.

As a consequence, if $\Phi = \mathcal{O}(V_m)$, the relative fitness - compared to the maximum attainable value - is $f = \frac{\Phi - \eta_d [M]^*}{\Phi} \approx \frac{V_m + (\eta_d - 1)K_M}{V_m + \eta_d K_M}$, which is not influenced by $\Phi$. As soon as $k_{cat}$ is sufficiently high, the level of flux does not impact the selective pressure acting on $k_f$ anymore but marginally, which helps understand the shape of the fitness landscape when an enzyme competes against degradation processes.

#### General description of nutrient driven pathways - numerical solution for the model with two enzymes

We checked the implementation of the Newton algorithm by comparing results for the flux of the second product given by the algorithm with those resulting from the numerical simulations of the process through Euler explicit method for 9 different couples of kinetic parameters, concentrated nearby the average enzyme kinetic parameters to avoid reactions too slow to reach equilibrium (low efficiencies) and/or numerical stability problems arising from high efficiencies. Simulations with the Euler method were ran for a total simulation time of 1s., with timesteps inversely proportional to the efficiency of the quickest kinetic parameter involved. Results for the control case of amino acids with a low degradation rate ($\eta_d = 10^{-4} s^{-1}$) and following a first enzyme relatively efficient ($k_f = 10^7 M^{-1} s^{-1}$ and $k_{cat} = 10^4 s^{-1}$) to avoid the aforementionned issues with stability are reported in the following tables:

| $log_{10} k_f$ \ $log_{10} k_{cat}$ | 2 | 3 | 4 |
|---|---|---|---|
| 5 | 9.08871051750e-07 | 9.09087090923e-07 | 9.09087909102e-07 |
| 6 | 9.09087909093e-07 | 9.09088727281e-07 | 9.09088809099e-07 |
| 7 | 9.09088809098e-07 | 9.09088890917e-07 | 9.09088899099e-07 |

Table S1: Flux at steady-state using Euler explicit method

| $log_{10} k_f$ \ $log_{10} k_{cat}$ | 2 | 3 | 4 |
|---|---|---|---|
| 5 | 9.09078909140e-07 | 9.09087090923e-07 | 9.09087909102e-07 |
| 6 | 9.09087909093e-07 | 9.09088727281e-07 | 9.09088809099e-07 |
| 7 | 9.09088809098e-07 | 9.09088890917e-07 | 9.09088899099e-07 |

Table S2: Flux at steady-state using Newton algorithm

Finally, we compared the relative difference between dynamical solutions using Euler method and the root finding Newton algorithm:

| $log_{10}k_{cat}$ / $log_{10}k_f$ | 2 | 3 | 4 |
|---|---|---|---|
| 5 | 0.000228646147092 | 6.40454889231e-13 | 2.40388743711e-13 |
| 6 | 3.4637409099e-13 | 2.14299849973e-14 | 1.44419451114e-14 |
| 7 | 2.05797717838e-13 | 1.9566504519e-14 | 4.30928964696e-15 |

Table S3: Relative differences between estimates of flux at steady-state

The relative difference between the estimates is very low except for $log_{10}k_{cat} = 2$ and $log_{10}k_f = 5$ because one second is not enough to reach steady-state for such kinetic parameters. Indeed, looking at the last two steps of dynamic simulation reveals that changes in flux cannot yet be overlooked in that case, as shown in this table where the difference corresponds to the one that would be observed if it were to remain constant during one second:

| $log_{10}k_{cat}$ / $log_{10}k_f$ | 2 | 3 | 4 |
|---|---|---|---|
| 5 | 1.74440671693e-09 | 0 | 0 |
| 6 | 0 | 0 | 0 |
| 7 | 0 | 0 | 0 |

Table S4: Difference of flux between the las two timesteps adjusted to one second

### Evolutionary trajectories for the simulations

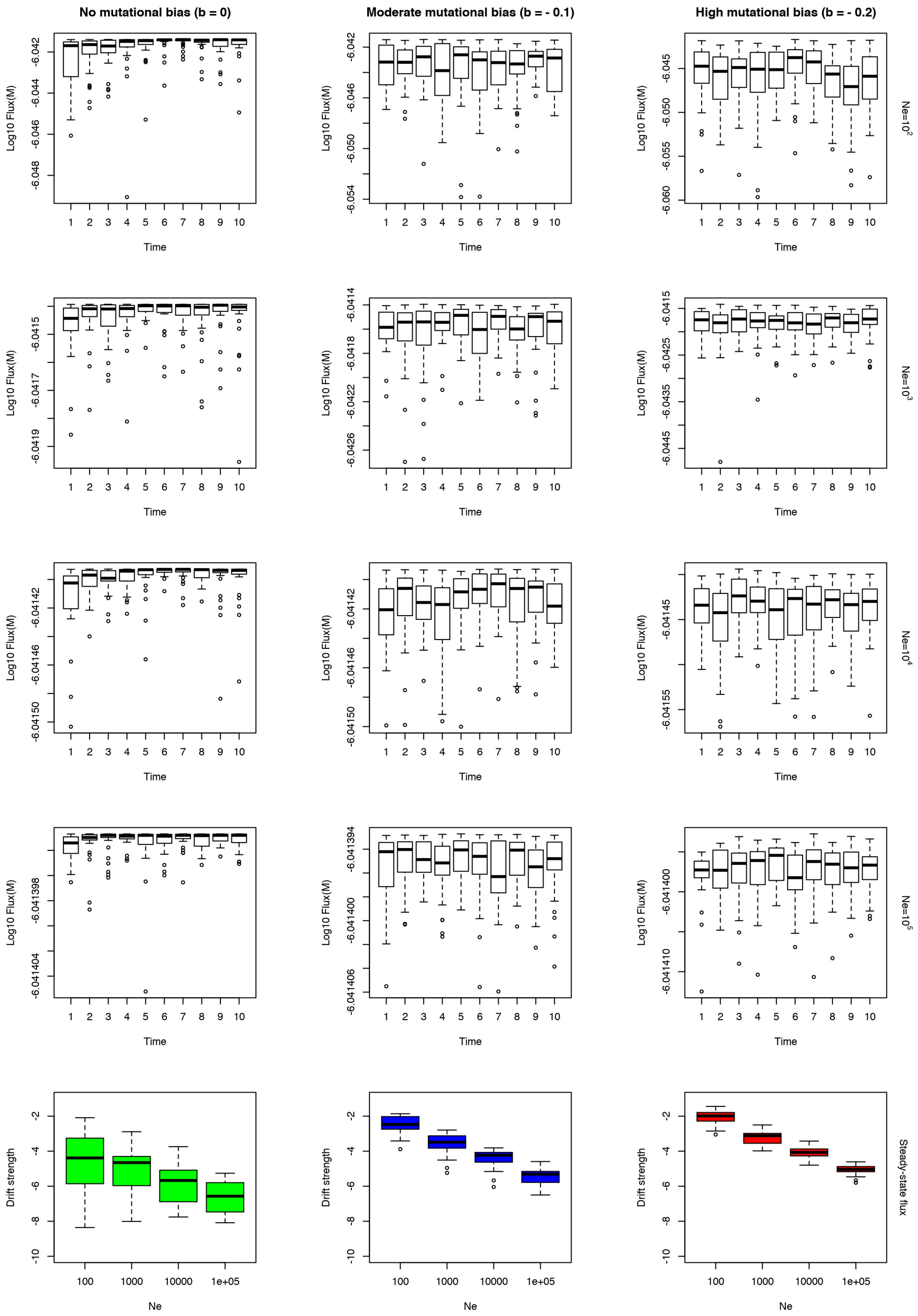


Figure S22: Evolutionary outcomes for enzyme fitnesses in the case of a moderate flux with high affinity ($V_{Tm} = 10^{-6}M$; $K_T = 10\mu M$). The first four lines represent the evolution of an enzyme's fitness relatively to the time, starting from an enzyme with a very low efficiency. Timesteps are specific to each set of parameters: they are proportional to $(2.5 \times 10^1)N_e$ for the low bias and increased by a factor 2 with a high bias and 4 with no bias for the establishment of steady-state is slowed down. Several parameters are considered : effective population sizes differ from one line to the other while mutational biases differ between columns. The last row indicates the span of fitnesses for each set of parameter when steady-state has been reached: each column corresponds to a different bias, as for the other plots.

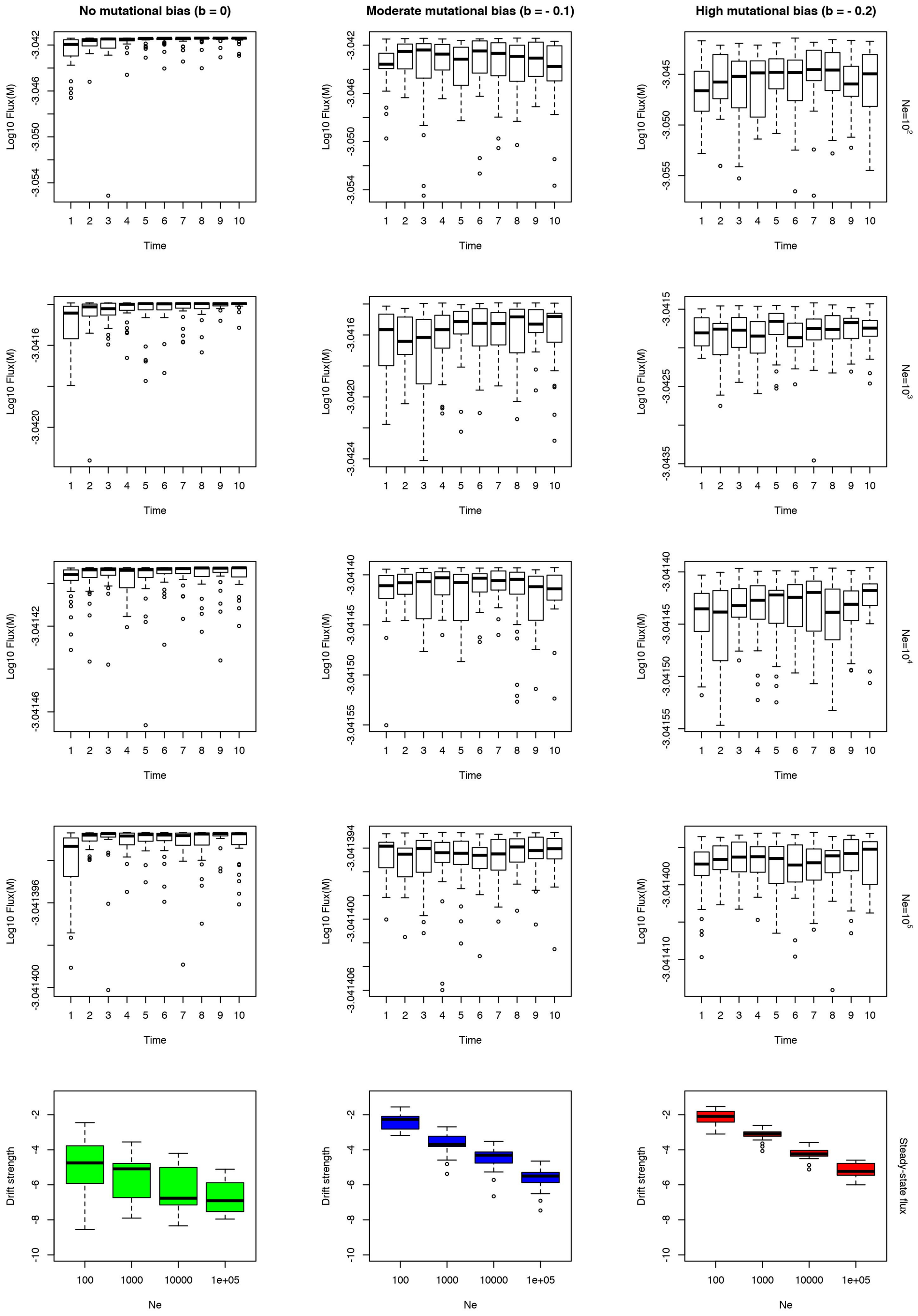


Figure S23: Evolutionary outcomes for enzyme fitnesses in the case of a high flux with low affinity ($V_{Tm} = 1mM$; $K_T = 0.1M$). The first four lines represent the evolution of an enzyme's fitness relatively to the time, starting from an enzyme with a very low efficiency. Timesteps are specific to each set of parameters: they are proportional to $(2.5 \times 10^1)N_e$ for the low bias and increased by a factor 2 with a high bias and 4 with no bias for the establishment of steady-state is slowed down. Several parameters are considered : effective population sizes differ from one line to the other while mutational biases differ between columns. The last row indicates the span of fitnesses for each set of parameter when steady-state has been reached: each column corresponds to a different bias, as for the other plots.

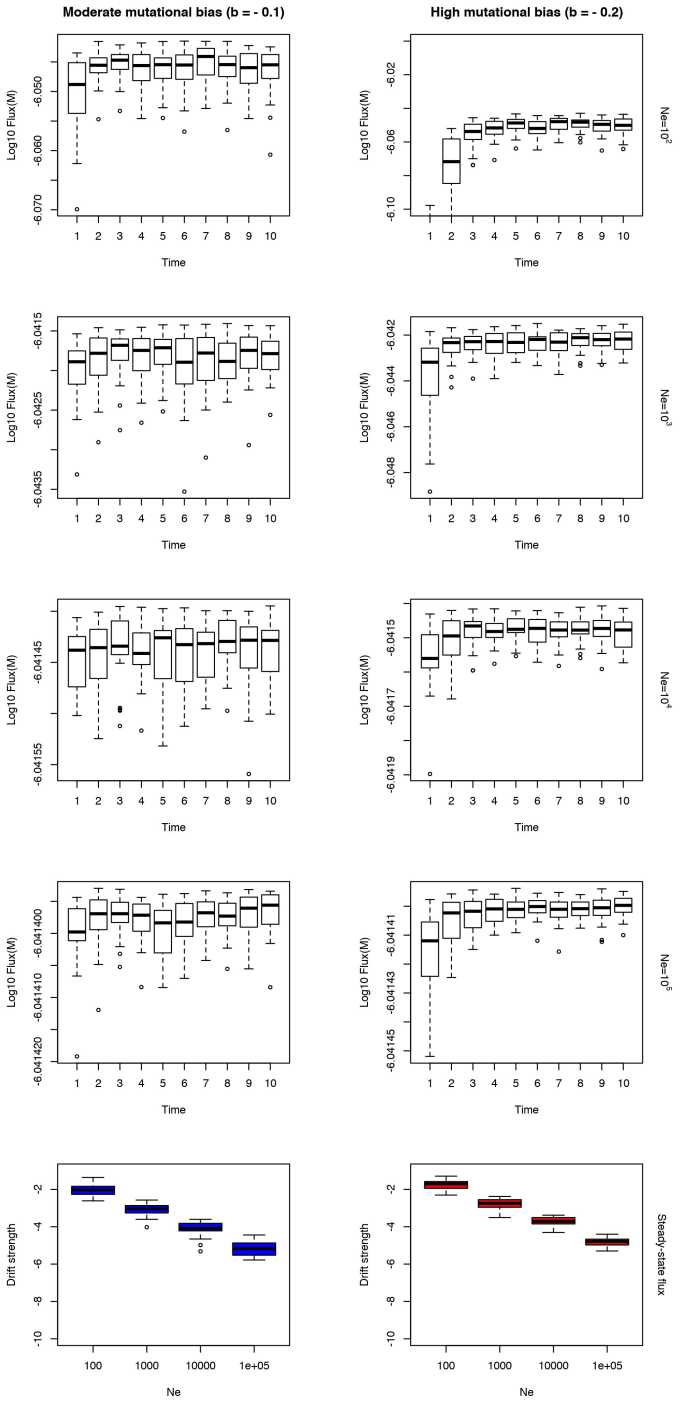


Figure S24: Evolutionary outcomes for enzyme fitnesses in the case of a moderate flux with high affinity ($V_{Tm} = 10^{-6}M$; $K_T = 10\mu M$) and a moderate trade-off ($\rho = -0.5$ between $k_{cat}$ and $k_f$). The first four lines represent the evolution of an enzyme's fitness relatively to the time, starting from an enzyme with a very low efficiency. Timesteps are specific to each set of parameters: they are proportional to $(10^2)N_e$ and increased by a factor 2 when mutational bias is increased for the establishment of steady-state is slowed down. Several parameters are considered : effective population sizes differ from one line to the other while mutational biases differ between columns. The last row indicates the span of fitnesses for each set of parameter when steady-state has been reached: each column corresponds to a different bias, as for the other plots.

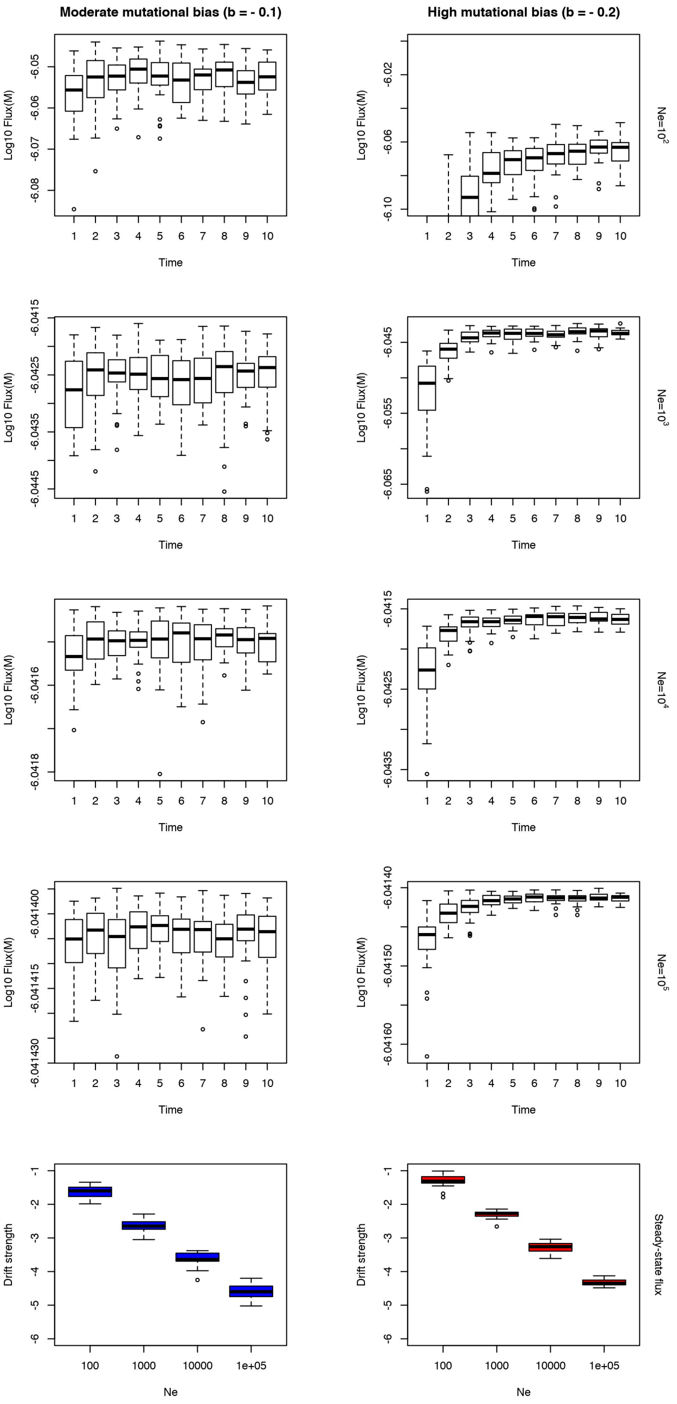


Figure S25: Evolutionary outcomes for enzyme fitnesses in the case of a moderate flux with high affinity ($V_{Tm} = 10^{-6}M$; $K_T = 10\mu M$) and a high trade-off ($\rho = -0.8$ between $k_{cat}$ and $k_f$). The first four lines represent the evolution of an enzyme's fitness relatively to the time, starting from an enzyme with a very low efficiency. Timesteps are specific to each set of parameters: they are proportional to $(2.5 \times 10^2)N_e$ and increased by a factor 2 when mutational bias is increased for the establishment of steady-state is slowed down. Several parameters are considered : effective population sizes differ from one line to the other while mutational biases differ between columns. The last row indicates the span of fitnesses for each set of parameter when steady-state has been reached : each column corresponds to a different bias, as for the other plots.

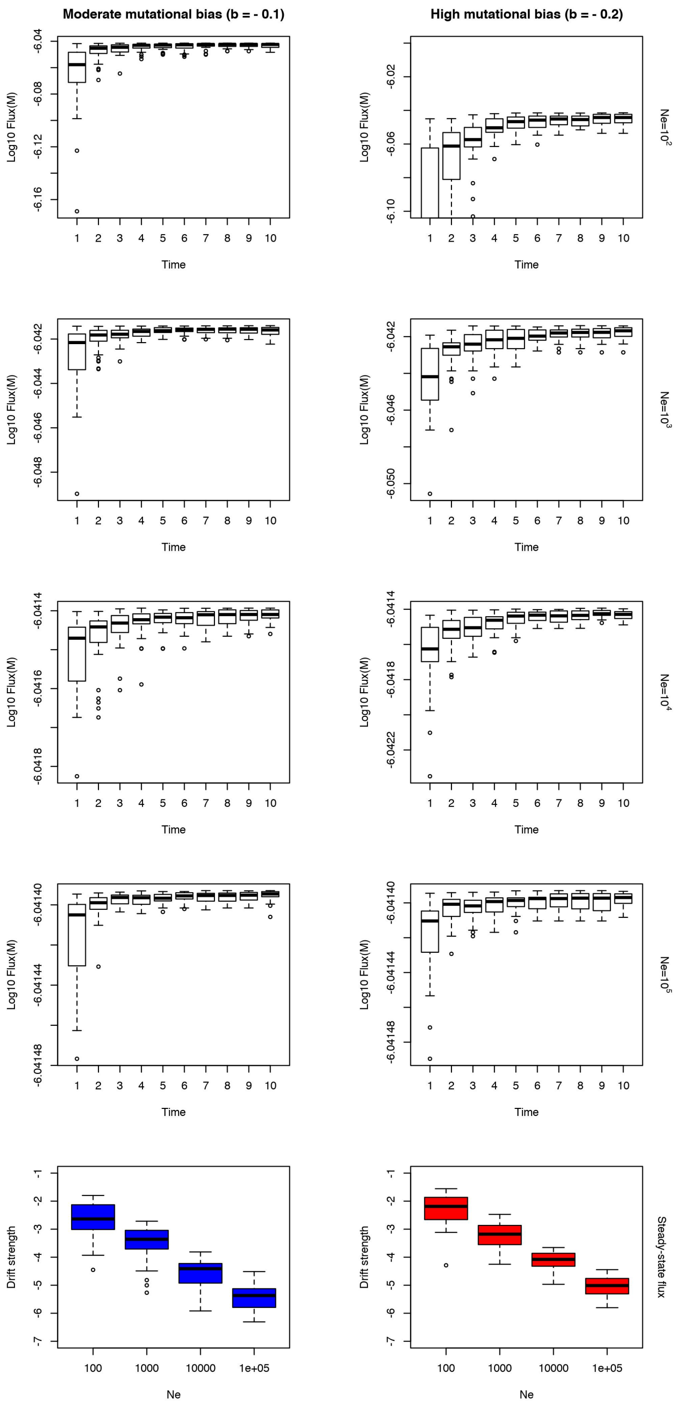


Figure S26: Evolutionary outcomes for enzyme fitnesses in the case of a moderate flux with high affinity ($V_{Tm} = 10^{-6}M$; $K_T = 10\mu M$) and a positive correlation ($\rho = 0.5$ between $k_{cat}$ and $k_f$) pushing both parameters in the same direction, for example if the respective energy barriers are positively correlated (which is unlikely, at least with this magnitude). The first four lines represent the evolution of an enzyme's fitness relatively to the time, starting from an enzyme with a very low efficiency. Timesteps are specific to each set of parameters: they are proportional to $(2.5 \times 10^1)N_e$ and increased by a factor 2 when mutational bias is increased for the establishment of steady-state is slowed down. Several parameters are considered : effective population sizes differ from one line to the other while mutational biases differ between columns. The last row indicates the span of fitnesses for each set of parameter when steady-state has been reached : each column corresponds to a different bias, as for the other plots.

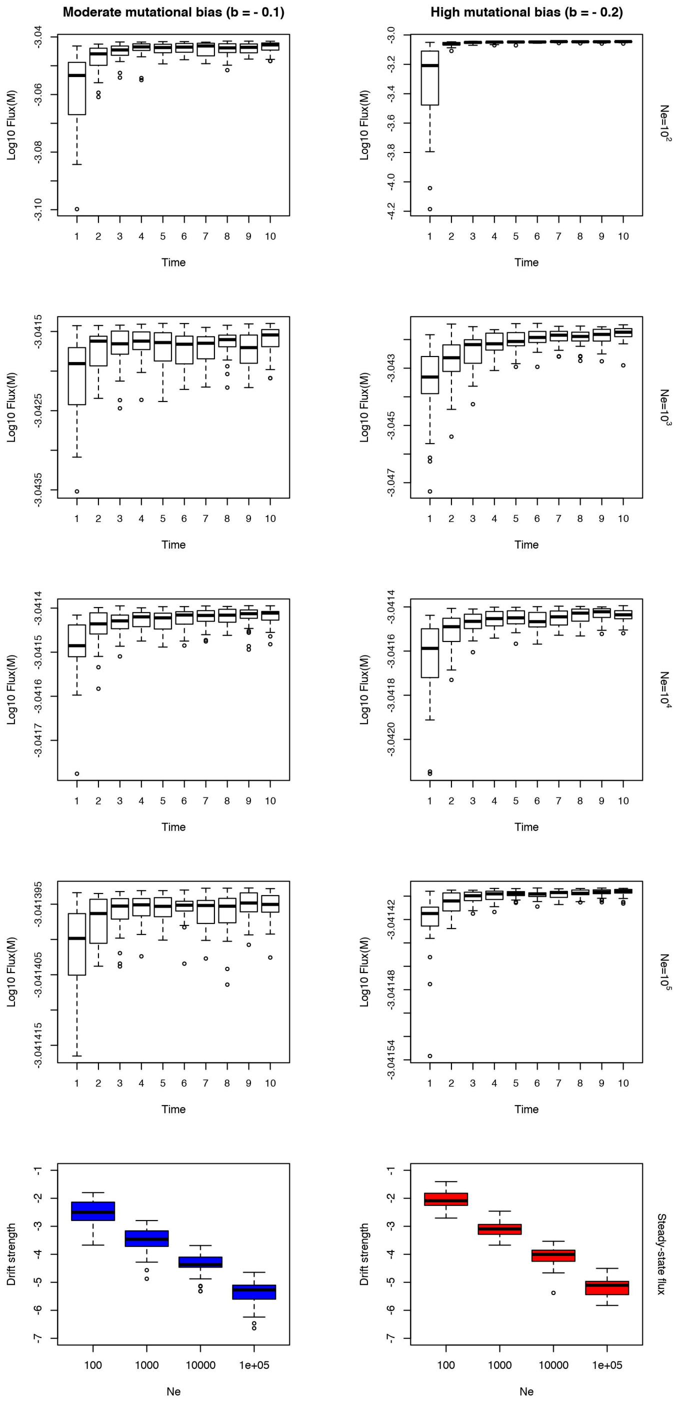


Figure S27: Evolutionary outcomes for enzyme fitnesses in the case of a high flux with low affinity ($V_{Tm} = 10^{-3}M$; $K_T = 1mM$) when expression is limited by macromolecular crowding effects. The first four lines represent the evolution of an enzyme's fitness relatively to the time, starting from an enzyme with a very low efficiency. Timesteps are specific to each set of parameters : they are proportional to $(2.5 \times 10^1)N_e$ and by a factor 2 when mutational bias is increased for the establishment of steady-state is slowed down. Several parameters are considered : effective population sizes differ from one line to the other while mutational biases differ between columns. The last row indicates the span of fitnesses for each set of parameter when steady-state has been reached : each column corresponds to a different bias, as for the other plots above.

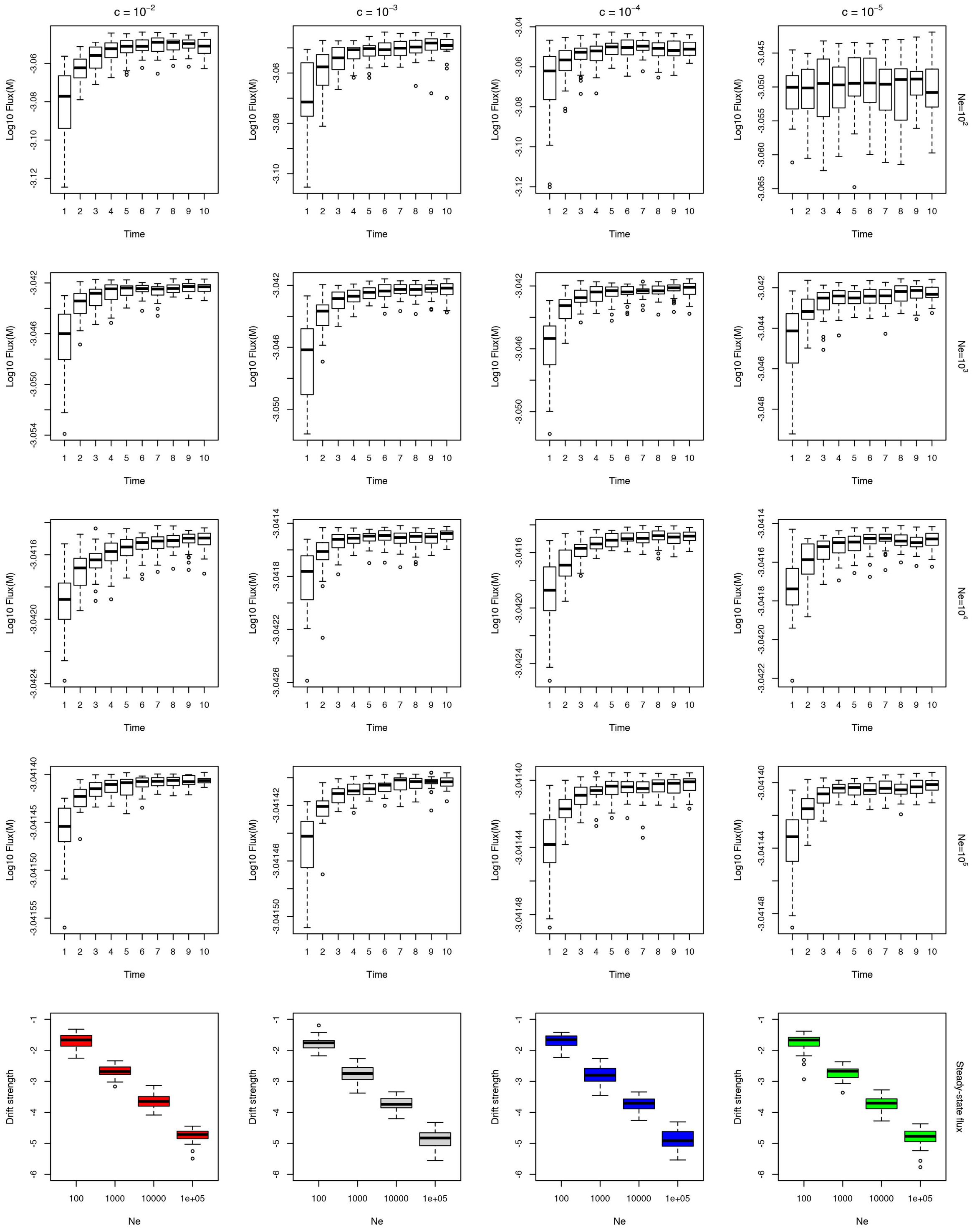


Figure S28: Evolutionary outcomes for enzyme fitnesses in the case of a high flux with low affinity ($V_{Tm} = 10^{-3}M$; $K_T = 1mM$) when expression is limited only by the cost of protein production. The first four lines represent the evolution of an enzyme's fitness relatively to the time, starting from an enzyme with a very low efficiency. Mutational bias was set to $b = -0.2$. Timesteps are specific to each set of parameters : they are proportional to $(5 \times 10^1)N_e$. Several parameters are considered : effective population sizes differ from one line to the other while costs of expression differ between columns. The last row indicates the span of fitnesses for each set of parameter when steady-state has been reached: each column corresponds to a different cost.

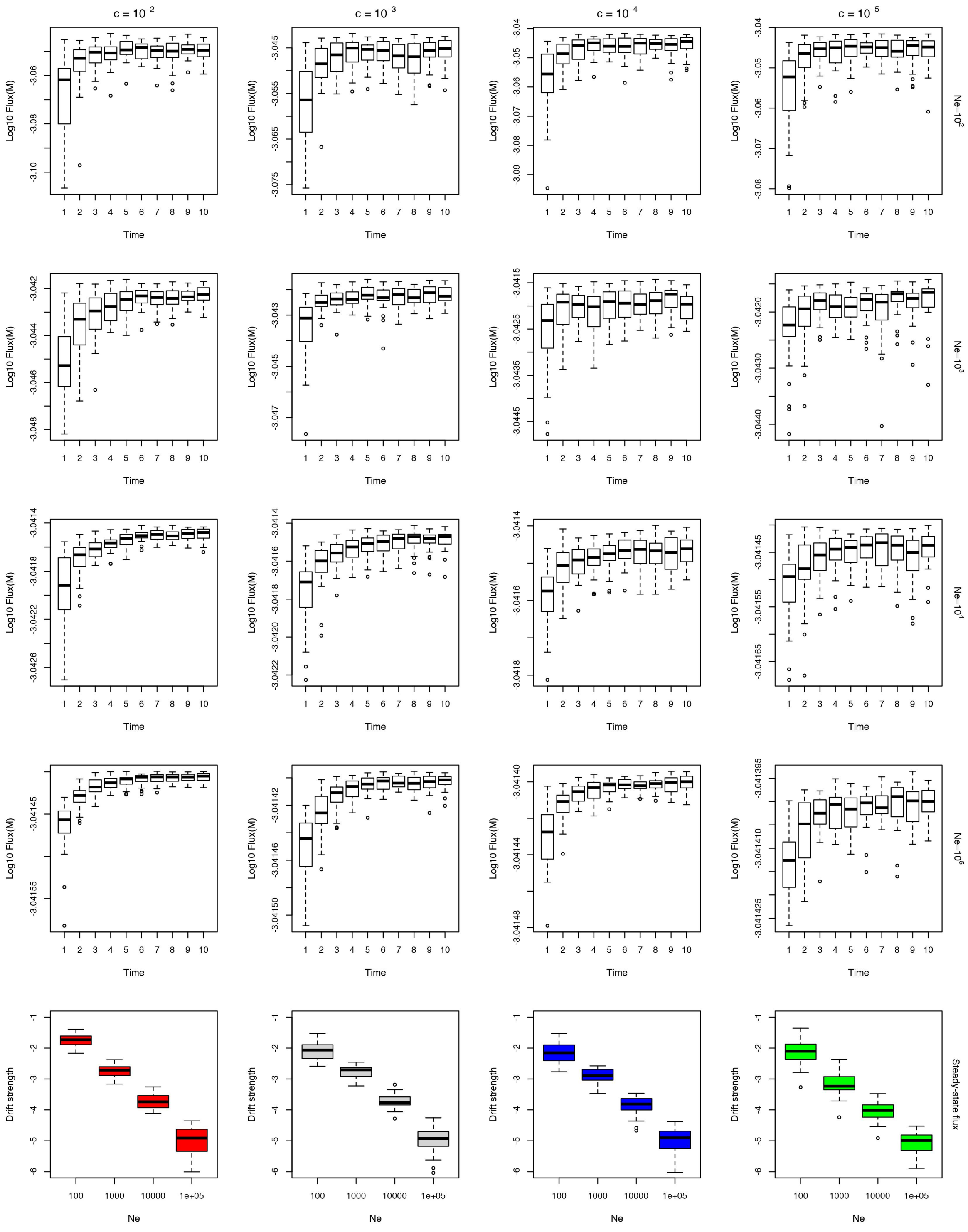

Figure S29: Evolutionary outcomes for enzyme fitnesses in the case of a high flux with low affinity ($V_{Tm} = 10^{-3}M$; $K_T = 1mM$) when the cost of expression includes both macromolecular crowding effects and the cost of production. The first four lines represent the evolution of an enzyme's fitness relatively to the time, starting from an enzyme with a very low efficiency. Mutational bias was set to $b = -0.2$. Timesteps are specific to each set of parameters : they are proportional to $(5 \times 10^1)N_e$. Several parameters are considered : effective population sizes differ from one line to the other while costs of expression differ between columns. Crowding influence is however considered fixed, since its effect should not be subject to a broad uncertainty/variability. The last row indicates the span of fitnesses for each set of parameter when steady-state has been reached : each column corresponds to a different cost, as for the other plots.
